# A deep learning-based approach to model anomalous diffusion of membrane proteins: The case of the nicotinic acetylcholine receptor

**DOI:** 10.1101/2021.08.16.456496

**Authors:** Héctor Buena Maizón, Francisco J. Barrantes

## Abstract

We present a concatenated deep-learning multiple neural network system for the analysis of single-molecule trajectories. We apply this machine learning-based analysis to characterize the translational diffusion of the nicotinic acetylcholine receptor at the plasma membrane, experimentally interrogated using superresolution optical microscopy. The receptor protein displays a heterogeneous diffusion behavior that goes beyond the ensemble level, with individual trajectories exhibiting more than one diffusive state, requiring the optimization of the neural networks through a hyperparameter analysis for different numbers of steps and durations, especially for short trajectories (<50 steps) where the accuracy of the models is most sensitive to localization errors. We next use the statistical models to test for Brownian, continuous-time random walk, and fractional Brownian motion, and introduce and implement an additional, two-state model combining Brownian walks and obstructed diffusion mechanisms, enabling us to partition the two-state trajectories into segments, each of which is independently subjected to multiple analysis. The concatenated multi-network system evaluates and selects those physical models that most accurately describe the receptor’s translational diffusion. We show that the two-state Brownian-obstructed diffusion model can account for the experimentally observed anomalous diffusion (mostly subdiffusive) of the population and the heterogeneous single-molecule behavior, accurately describing the majority (72.5% to 88.7% for α-bungarotoxin-labeled receptor and between 73.5% and 90.3% for antibody-labeled molecules) of the experimentally observed trajectories, with only ∼15% of the trajectories fitting to the fractional Brownian motion model.

## Introduction

The nicotinic acetylcholine receptor (nAChR) is the archetype member of the superfamily of pentameric ligand-gated ion channels (see (Changeux, 2018) for a review). The pentameric nAChR intervenes in a variety of physiological processes and is subject to dysfunction in a variety of diseases of the central nervous system such as Alzheimer and Parkinson diseases, schizophrenia spectrum disorders, autistic spectrum disorders, as well as in diseases of the peripheral nervous system like myasthenia gravis (Vincent, 2002;Paz and Barrantes, 2019;2021).

In myasthenia gravis, the muscle-type nAChR is degraded after autoimmune attack as a result of autoantibody binding to the receptor and its subsequent complement-mediated internalization and destruction of the protein. We have recently applied stochastic optical reconstruction microscopy (STORM) (Bates et al., 2013) in its direct mode (dSTORM) (Andronov et al., 2021), a form of single-molecule localization microscopy (SMLM) techniques (Deutsch et al., 2012;Nair et al., 2013;Manzo and Garcia-Parajo, 2015), to study the dynamics of the receptor cross-linked with a multivalent monoclonal antibody. The nAChR trajectories were followed with the single-particle tracking technique (Alcor et al., 2009). Customarily, dSTORM is intended for fixed samples. We have optimized image acquisition protocols using this technique on live cells for short times (Mosqueira et al., 2018;Mosqueira et al., 2020). The acquired trajectories with short durations (usually < 50 steps) pose inherent challenges such as sensitivity to the localization error of methods such as mean-squared displacement (MSD).

In recent years, artificial intelligence and in particular its subset, machine learning (ML) has had an important impact on microscopy, especially in cryo-electron microscopy and optical superresolution microcopy (nanoscopy). One of the ML methodologies proving to be ground-breaking is the application of deep neural networks for analyzing superresolution microscopy data. This is due to their capacity to automatically detect singular objects or complex patterns, outperforming the more traditional algorithms for image processing owing to their much-reduced computation time, thus facilitating the evaluation of vast amounts of experimental data in relatively short times.

The analysis of trajectories, a sequence of particle (molecule) locations sampled over time, and more generally, of time-series data using deep learning, can be approached with different architectures, e.g., recurrent neural networks (Yu et al., 2019), fully convolutional networks (Wang et al., 2017), graph neural networks (Scarselli et al., 2009) or residual neural networks (He et al., 2016), among others. In all cases, the actual performance of the selected approach will determine the most suitable architecture. In this work we have trained and optimized several deep neural networks with simulated data (on average 50,000-60,000 tracks) that emulate different modalities of protein diffusion in membranes (Krapf, 2015). Single-molecule experimental data obtained with the STORM technique were then analyzed with these trained networks to find the physical models that best describe the diffusional properties of the observed trajectories. These models are the fractional Brownian motion (fBm), continuous time random walk, two physical models previously applied in the work by Shechtman and coworkers (Granik et al., 2019). We introduce a set of modifications to the neural networks proposed by these authors and by Volpe and coworkers (Bo et al., 2019) to characterize the diffusion properties at the single-molecule level, and we additionally introduce a two-state model which combines Brownian and obstructed diffusion based on the switching diffusion concept (Grebenkov, 2019). Previous results from our laboratory analyzed using classical methods such as mean squared displacement analysis (MSD) suggested that this model could explain the anomalous diffusion behavior of some of the nAChR trajectories (Mosqueira et al., 2018;Mosqueira et al., 2020). To the best of our knowledge, the analysis of the nAChR dynamics utilizing neural networks presents an original use of machine learning,

## Results

### STORM imaging of nicotinic acetylcholine receptor molecules in live cells

The dynamics of nAChR molecules were interrogated with nanoscopy in live cells using a heterologous mammalian cell expression system, the CHO-K1/A5 clonal cell line (Roccamo et al., 1999). These cells robustly express the muscle-type receptor at its cell surface. Cells were labeled with either a nicotinic antagonist, the small peptide α-bungarotoxin (BTX) tagged with Alexa Fluor^555^, or the monoclonal antibody mAb35 (mAb) followed by a secondary antibody labeled with Texas Red. Sets of cells were treated with methyl-β-cyclodextrin (CDx) for acute depletion of the cell-surface cholesterol content, or with CDx-cholesterol (CDx-Chol) complexes to enrich membranes with the sterol (Borroni et al., 2007).

Single-particle tracking superresolution microscopy provided sets of data in the form of successive micrographs (a “stream”) of selected areas of different cells acquired at a rate of 10 ms/frame. The time-dependent localization of the molecules and their trajectories in the plane of the membrane were reconstructed as detailed in Material and Methods. We imaged the flat, ventral membrane of the cell adhered to the coverslip (Figure 1).

**Figure 1.**
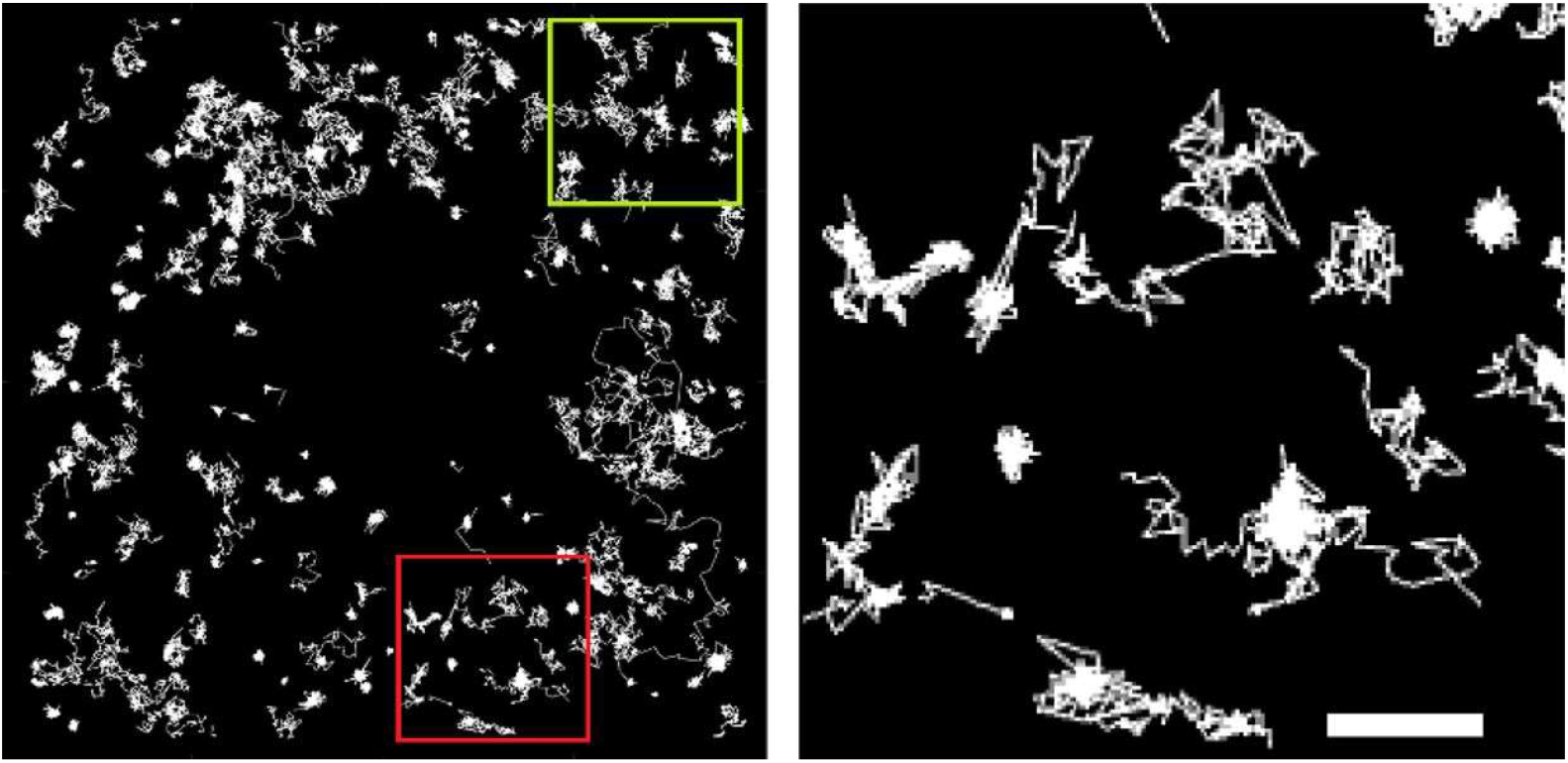
Representative reconstructed BTX-labeled nAChR trajectories in a 10 × 10 μm square region of a CHO-K1/A5 cell (left panel) is magnified on the right panel to facilitate observation of trajectories with free Brownian walks and stopovers, and other essentially immobile single-molecule particles. See Suppl. Material for details. The bar on the right panel corresponds to 500 nm.

Prior to submitting the experimental data to the networks, the procedure of Golan and Sherman (Golan and Sherman, 2017) was applied to the reconstructed sets of localizations. This procedure filters out trajectories which, based on the radius of gyration and the mean step size of the particle’s displacement (Suppl. Material Eq. 3), can be considered immobile (stationary). Approximately 60.5% and 76.3% of the total single-molecule trajectories were considered immobile for BTX- and mAb-labeled samples, respectively.

### Deep neural network classification of mobile nAChR trajectories: initial physical model

Deep neural networks trained with simulated data as described in the Supplementary Material and Methods section were applied to the mobile single-molecule trajectories. To find the physical model best able to describe the behavior of the nAChR lateral motion in both BTX- and mAb-labeled samples under control and cholesterol-modified conditions, the Physical Model classification network of Figure 2 was applied first. The network architecture is based on the characterization network introduced by Shechtman and coworkers (Granik et al., 2019) which relies on the concept of Temporal Convolutional Networks (Bai et al., 2018). We introduce here a new model, based on the 2-state model of Grebenkov (Grebenkov, 2019), combining a switching behavior of the diffusing particle between Brownian diffusion and obstructed diffusion. The new model trajectories are simulated covering a broad range of values for the diffusion coefficient, switching rate for each individual state (Supplementary Table 1), and area of confinement of the obstructed diffusion state (see Suppl. Material). The hyper-parameters were fine-tuned to obtain the best performance for a varied range of trajectories with different time duration and number of steps. Models for evaluating trajectories with less than 50 steps require precise optimization for setting hyperparameters able to correctly generalize in this range and to produce the highest possible accuracy of the model.

**Figure 2.**
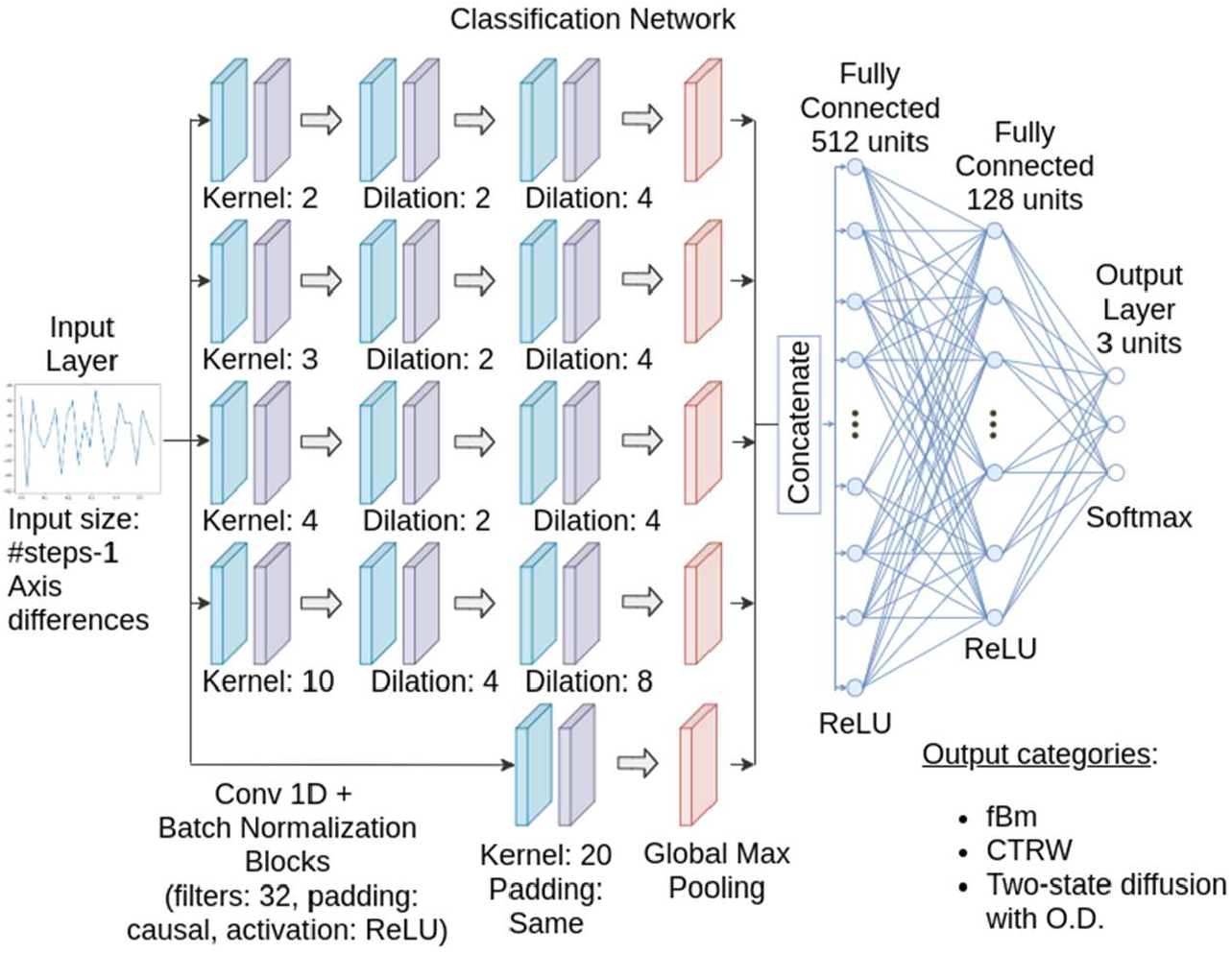
The Physical Model classification network receives as input all the validated mobile trajectories after the Golan and Sherman filtering criterion (Golan and Sherman, 2017) within the range of 25-900 steps. The category with the highest value is selected as the output. This classification is key to the analysis, separating the trajectories for the fBm classification network and two-state segmentation network. The architecture is based on the methods introduced by Shechtman and coworkers (Granik et al., 2019) which rely on the concept of TCN (Bai et al., 2018).

The Physical Model network (Figure 2) was trained to classify trajectories according to three different physical models: the continuous time random walk (CTRW) model, the fractional Brownian motion (fBm) model and the two-state (Brownian diffusion + obstructed diffusion) model. The network receives a zero-centered vector with the n-th discrete difference between positions along a given axis. The output assigns a probability for each category, and the one with the highest value is chosen.

As shown in Figure 3, the two-state model appears to account for most of the mobile trajectories under all experimental conditions tested. The proportion of trajectories that satisfactorily fitted the 2-state model is higher for the mAb-labeled samples. In the case of BTX, this model accounts for 72.5% to 88.7%, and for mAb, between 73.5% and 90.3% of the trajectories. The actual percentages are listed in Supplementary Table 2.

**Figure 3.**
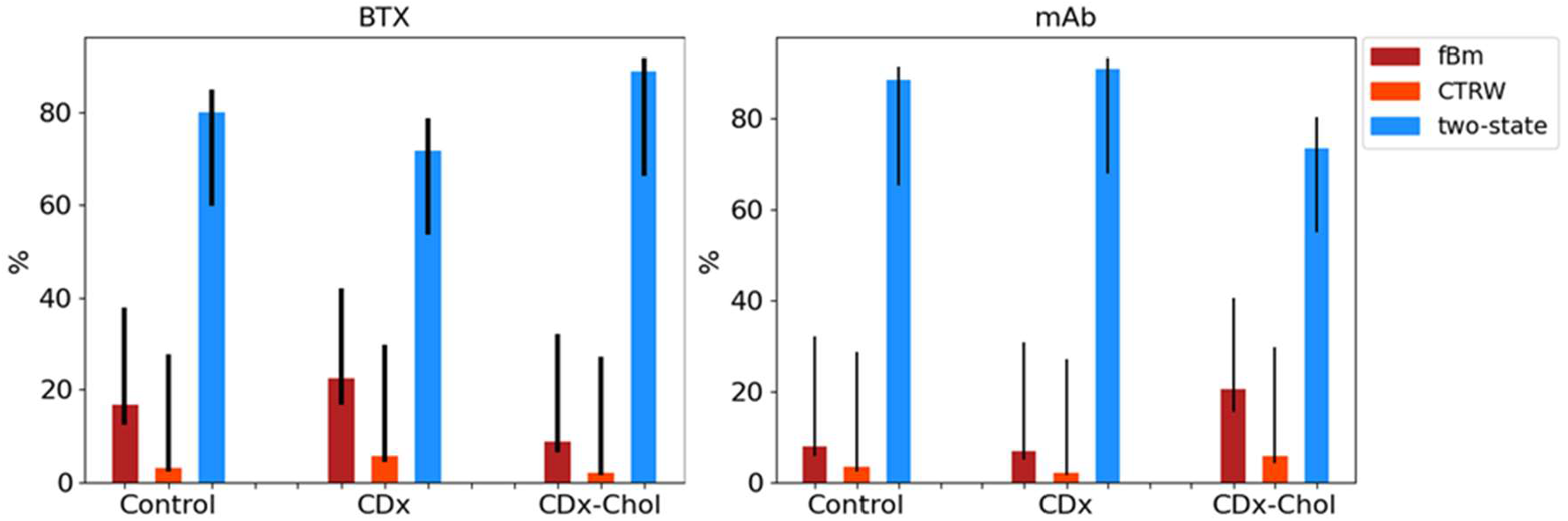
The Physical Model classification network of Figure 2 was applied to the complete dataset of nAChR mobile trajectories (25 to 900 steps) to yield the most probable physical model of diffusion. Three models: fractional Brownian motion (fBM), continuous-time random walk (CTRW), and two-state (Brownian diffusion + obstructed diffusion) models were tested. As can be appreciated, the two-state model appears to be the most likely representation of the mobile trajectories. The black lines indicate the range of values below the 95% confidence interval of the classification error of all the trained models of the classification network.

### The fBm covers a wide range of motional regimes

Those trajectories previously characterized by the Three Physical Model classification network (Figure 2) as fBm, were further analyzed by the fBm Subclassification network (Figure 4). Simulated trajectories conforming to the fBm model but with three ranges of Hurst indices (Mandelbrot and Van Ness, 1968) were fed to the network and used in the training process. The output of the fBm Subclassification network separated the trajectories into three categories of motional regimes: subdiffusive, Brownian and superdiffusive. The network architecture shares similarities with the Physical Model network of Figure 2; however, more convolutional blocks were added and the number of units in the fully connected layers were incremented to improve performance with respect to the original architecture. The reason for this choice lies in the complexity of classifying short trajectories with high accuracy. The network receives a zero-centered vector with the n-th discrete difference between positions along a given axis. The output assigns a probability for each category, and the one with the highest value is chosen.

**Figure 4.**
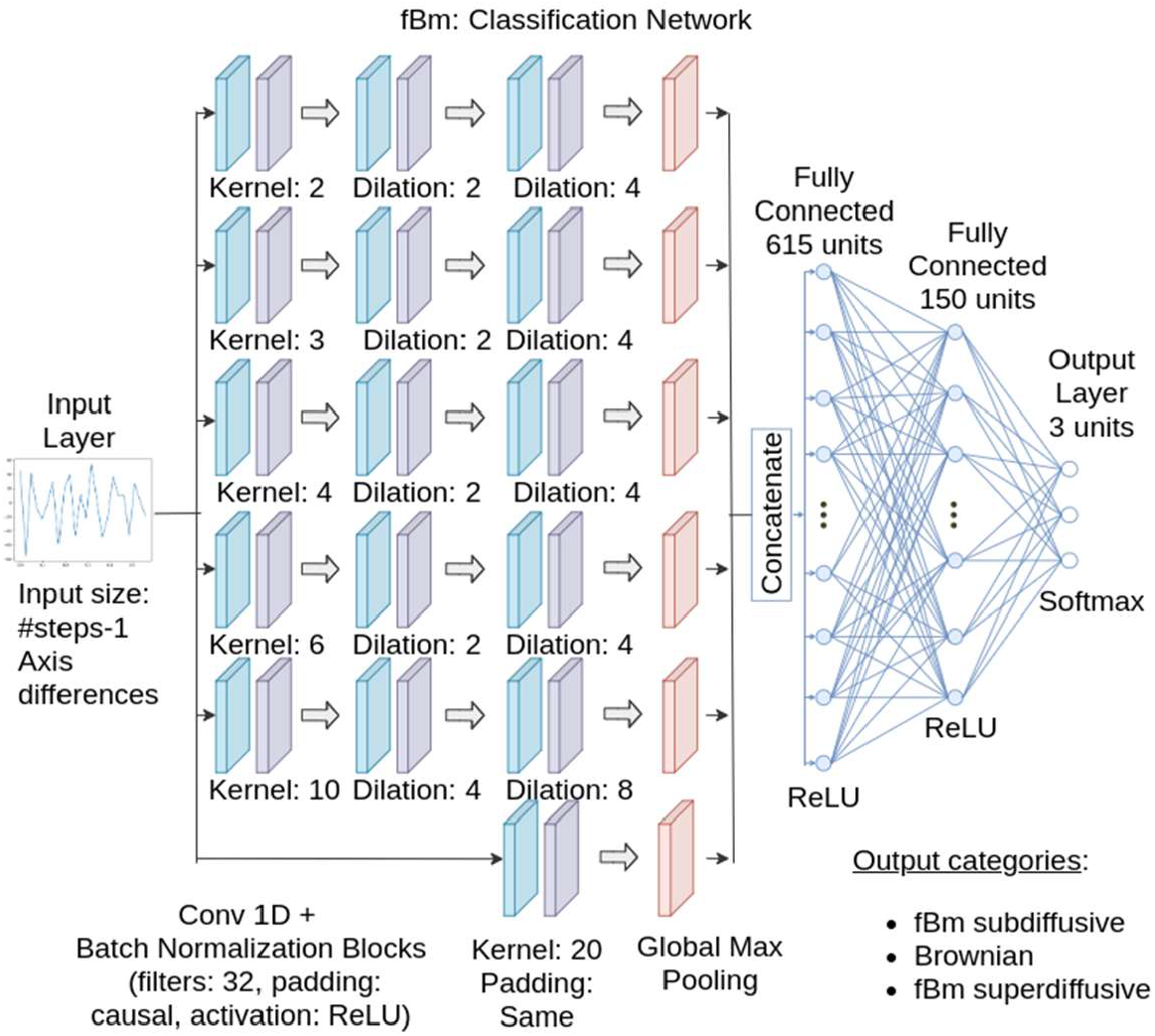
Diagram showing the fBm Subclassification network, designed to subcategorize the motional regime of the nAChR trajectories previously classified as fBm. Each category (motional regime) represents a range of values of the Hurst Index for subdiffusive [0.1-0.42], Brownian [0.43-0.57] and superdiffusive [0.58-0.9] motion. This network not only yields the predominant type of fBm diffusion but also information to select a more specific Hurst Exponent Network, trained specifically for a smaller range of data.

The result of the output layer yielded the probability for each category. The results (Figure 5) show that most of the trajectories exhibiting fractional Brownian motion are predominantly subdiffusive for the three experimental conditions (control, cholesterol-depleted (CDx) and cholesterol-enriched (CDx-Chol) and for the two labeling protocols (BTX and mAb), the rest displaying Brownian motion (11.3-25%) and a negligible proportion of superdiffusive trajectories. The numerical results are presented in Supplementary Table 3.

**Figure 5.**
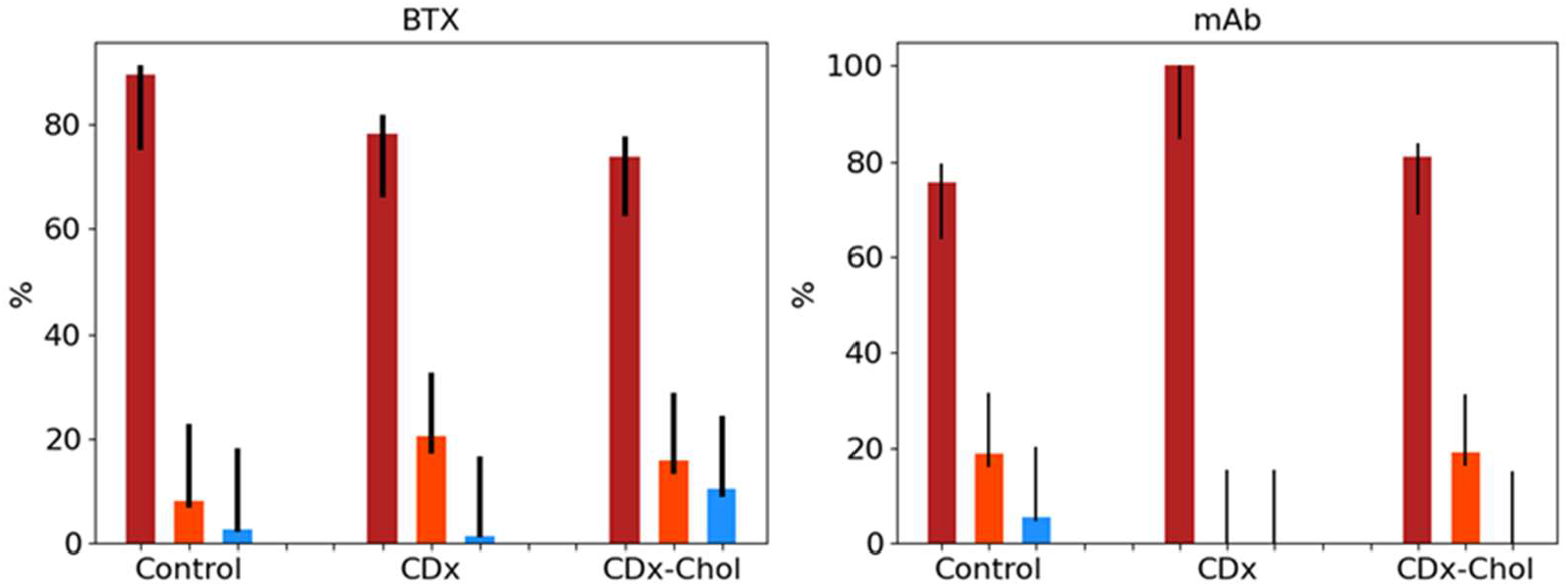
The output of the fBm Subclassification network of Figure 4 categorizes trajectories into subdiffusive (burgundy), Brownian (red), and superdiffusive (blue) motion. The input trajectories covered the entire range (25 to 900 steps) of the data previously classified as fBm using the network of Figure 2. The black lines indicate the range of values below the 95% confidence interval of the classification error of all the trained models of the classification network.

### Hurst Exponent network

All the trajectories classified as fBm and further subcategorized into subdiffusive, Brownian, and superdiffusive were fed to the Hurst Exponent Network (Suppl. Figure 2).

In the case of fractional Brownian motion, the anomalous exponent (see Suppl Material Eq. 4) is frequently parameterized with the Hurst (Mandelbrot and Van Ness, 1968) or self-similarity parameter. These exponents are linked by the following relation: 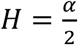. The Hurst exponent (Mandelbrot and Van Ness, 1968)measures how quickly a particle diffuses, quantifying whether the trajectory will return to a mean position, or diffuse in another direction. The first condition is known as subdifussion and covers Hurst indices in the range [0, 0.5). The second condition refers to a superdiffusive regime with Hurst exponent values greater than 0.5. The H = 0.5 is a special case, the classical Brownian motion. As discussed in the literature (see reviews in (Krapf, 2015;2018)), the viscoelastic properties of the plasma membrane may provide the ideal conditions for membrane-embedded proteins to exhibit fractional Brownian motion type of diffusion.

The Long-Short Term Memory architecture introduced by Volpe and coworkers (Bo et al., 2019), based on the Long-Short term memory neural networks (Hochreiter and Schmidhuber, 1997), has been used to analyze the anomalous diffusion coefficient that predicts the value of the anomalous exponent. We used this approach here to predict the value of the Hurst Index (Mandelbrot and Van Ness, 1968). The fBm category was used as prior information to select a model specified for the diffusion range. This network (Suppl. Figure 2) receives a 2-dimensional vector with each localization coordinate (normalized to zero-mean and unitary standard deviation), and the time vector adjusted to [0, 1]. The output is the Hurst Index. Additionally, we introduced a fully connected layer with a 128 unit-SELU activation function (Klambauer et al., 2017) to the output layer. As shown in Suppl. Figure 10, this allowed us to obtain more accurate results than the original work. The Hurst exponent mean, 95% CI of the mean, and 95% CI are shown in Figure 6. No statistically significant differences were found between the experimental conditions and labels. The numerical results are presented in Supplementary Table 6.

**Figure 6.**
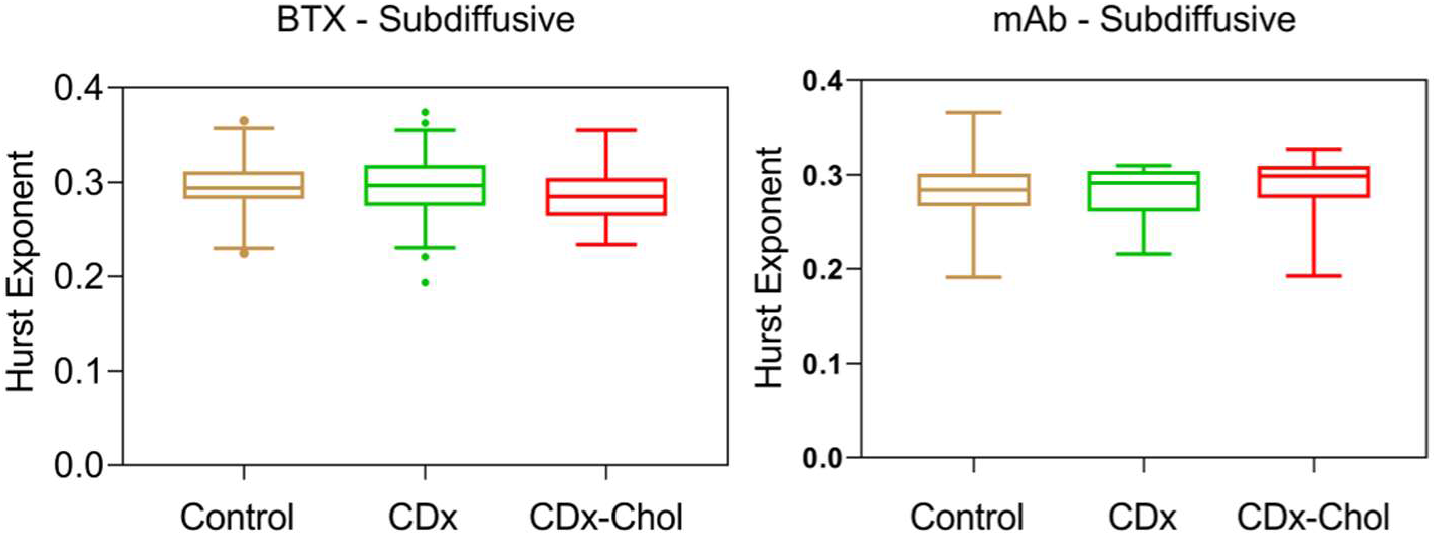
Hurst exponent values. As can be seen in both BTX- and mAb-labelled samples, there are no statistically significant differences between the three datasets under the three experimental conditions. Also, no statistically significant differences were found between BTX and mAb samples. Whiskers in box plots correspond to the interquartile range; the limits indicate 2.5^th^ and 97.5^th^ percentiles; the horizontal lines are the median in each case.

### The two-state model: confined and free-diffusion portions

The great majority of the nAChR trajectories appear to conform to the two-state physical model (Figure 3). Previous work from our laboratory showed that free-diffusing individual nAChR trajectories were interrupted by confinement sojourns of a few hundred millisecond duration (Mosqueira et al., 2018;Mosqueira et al., 2020). Turning-angle analysis (Burov et al., 2013;Sadegh et al., 2017) and escape-time analysis (Weigel et al., 2011) led us to conclude that the hindrance to motion experienced by diffusing molecules in the confined state was due to physical impediments, and the steepness of the turning-angle curves indicated high density of these physical obstacles, in agreement with an obstructed diffusion case (Weigel et al., 2012;Burov et al., 2013) (reviewed in (Krapf, 2015)).

Trajectories classified as two-state by the initial network (Figure 2) were further analyzed by the Segmentation network (Figure 7) to discriminate between the two individual states. This network is based on Temporal Convolutional Networks (Bai et al., 2018) but there are some differences between the two classification networks: the input in our network considers the position at each step, and not the difference between two consecutive steps as in the Physical Model classification network (Figure 2). Additionally, the pooling after the convolutional blocks is over the average of the multiple convolutions, and the number of units in the last dense layers is directly related to the number of steps in the trajectory.

**Figure 7.**
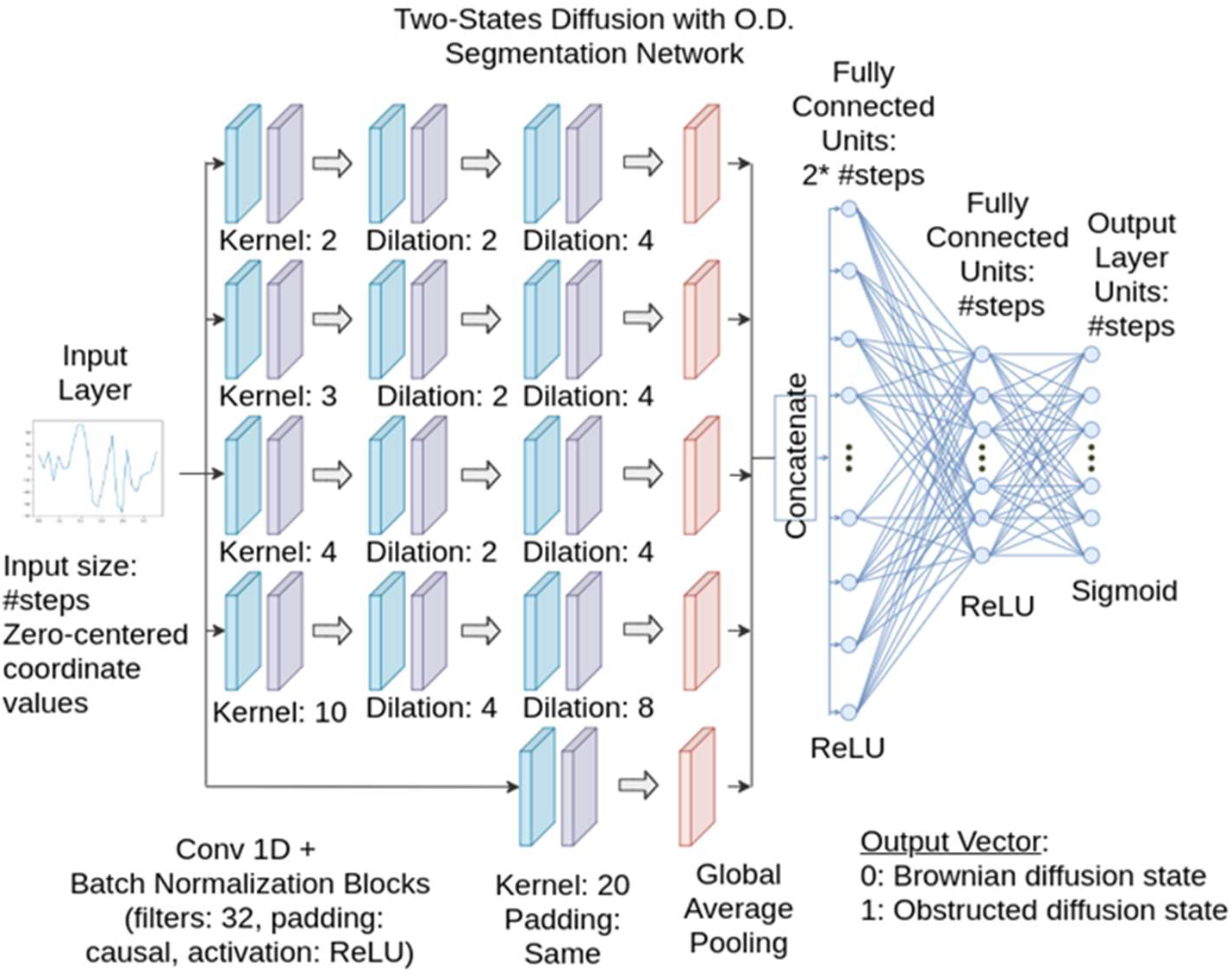
The Segmentation Network of the two-state model receives as input the trajectories classified as belonging to the 2-state model by the Three Physical Model network of Figure 2. The architecture was designed using the strategy of Shechtman and coworkers (Granik et al., 2019), with changes introduced in the input, pooling, and output layers in our version of the network to accomplish new tasks: i) the input considers the position at each step, unlike other conditions where the variation of the process over time is relevant; ii) the pooling is done over an average of the convolutions, providing better results than those obtained with a max pooling procedure, and iii) the dense layers reduce the size of the units by a factor of two at each level and the output layer has an output unit for every step.

The network is trained to classify each step and select the most probable state, either 0 or 1, zero corresponding to Brownian diffusion, and 1 to the alternate obstructed diffusion state. The network receives a 1-dimensional vector, with the position at each step normalized to a zero-mean. The output is also a 1-dimensional vector with the most probable state at each step.

Next, this information was used to compute the residence times of the free-diffusing (Brownian) and confined states of the trajectories and the confinement area in the obstructed diffusion state, respectively, as listed in Table 1. The residence time is defined as the duration (measured in seconds) of a segment belonging to the Brownian or obstructed diffusion state, respectively, in other words the difference in the time vector between the last and first localization of each segment. After evaluating the trajectories classified as two-state by the classification network together with the segmentation network, a filter was applied to remove the segments smaller than 3 steps, to avoid analyzing segments incorrectly classified to one of the possible states. The value for the filter was selected considering independence between each output and selecting a probability of misclassification of 0.01. Additionally, in some cases (∼14%) the segmentation was not able to detect more than one segment (single state trajectory, Brownian in all cases), due to e.g. the inability of the network to detect correctly or to the misclassification of truly Brownian trajectories as two-state ones.

**Table 1.** Hypoexponential fitting (Suppl. Material Eq. 9) of the residence time (expressed in seconds) in the Brownian diffusion state and obstructed diffusion state of the two-state model following classification of the data using the Segmentation network of Fig. 5. In Supplementary Figures 3 and 4, a graphical depiction of these results is shown in the form of the empirical probability density function and the hypoexponential fitting, which is a function of the residence time *τ*_1_ and *τ*_2_.

|  |  | Brownian diffusion state |  |  |
| --- | --- | --- | --- | --- |
|  |  | Control | CDx | CDx-Chol |
| BTX | $\tau_1$ (s) | $0.358 \pm 0.004$ | $0.279 \pm 0.004$ | $0.287 \pm 0.005$ |
| | $\tau_2$ (s) | $0.025 \pm 0.001$ | $0.024 \pm 0.001$ | $0.077 \pm 0.003$ |
| mAb | $\tau_1$ (s) | $0.202 \pm 0.006$ | $0.228 \pm 0.003$ | $0.293 \pm 0.003$ |
| | $\tau_2$ (s) | $0.066 \pm 0.003$ | $0.02 \pm 0.001$ | $0.023 \pm 0.001$ |
|  |  | Obstructed diffusion state |  |  |
| BTX | $\tau_1$ (s) | $0.051 \pm 0.002$ | $0.075 \pm 0.003$ | $0.082 \pm 0.002$ |
| | $\tau_2$ (s) | $0.017 \pm 0.001$ | $0.015 \pm 0.001$ | $0.016 \pm 0.001$ |
| mAb | $\tau_1$ (s) | $0.111 \pm 0.023$ | $0.146 \pm 0.003$ | $0.123 \pm 0.002$ |
| | $\tau_2$ (s) | $0.032 \pm 0.012$ | $0.027 \pm 0.001$ | $0.013 \pm 0.001$ |

The probability density function of the residence times for the different experimental conditions in the free-diffusing Brownian state and the obstructed diffusion state are shown in Suppl. Figures 3 and 4, respectively.

Comparison of the results observed under the different experimental conditions (control, cholesterol depletion (CDx), and cholesterol enrichment (CDx-Chol)) in the Brownian state showed statistically significant differences between BTX control and BTX CDx (p < 0.05). As shown in Table 1, the longer mean residence time for the cholesterol depletion condition (0.28 s) is shorter than the lifetime in the control (0.36 s). When BTX CDx was compared with CDx-Chol, statistically significant differences were apparent (p < 0.01). The shorter mean residence time increased upon cholesterol enrichment (CDx-Chol). In the case of mAb, changes in cholesterol content did not show statistically significant differences. Statistical differences were very pronounced (p < 0.0001) between a given mAb or BTX condition and its homologous condition (e.g. BTX control vs. mAb control). This is valid for all experimental conditions. In general, mAb-tagged receptors exhibit smaller residence times for longer segments, except for CDx-Chol.

We performed similar comparisons for the obstructed diffusion state. The results showed significant differences between BTX control vs. the two cholesterol-modifying conditions (p < 0.0001). In both cases, the shorter mean residence time increased relative to the control condition. When the two labeling procedures were contrasted, we found that BTX labeled-vs. mAb labeled-samples also exhibited highly significant statistical differences (p < 0.0001). For the shorter mean residence time, BTX-tagged nAChR trajectories exhibited shorter times than mAb under control and CDx conditions, and the longer residence times observed in longer segments were longer in mAb-labeled samples.

We also analyzed the areas covered by the confined state portion of the trajectories (Figure 8). The confinement area is defined as the difference between the maximum and minimum value in each coordinate of the localizations for each segment. In general, BTX showed smaller areas than mAb, with values between 0.0014 to 0.0019 versus 0.0024 to 0.0028 µm^2^. We found statistical differences in the distribution of the data between the different experimental conditions for mAb-vs. BTX-labeled samples, varying from p < 0.0001, p < 0.05, and p < 0.1 for control, CDx, and CDx-Chol, respectively. No statistical differences were found between the different experimental conditions labeled with the same probe. Numerical values are listed in Suppl. Table 5.

**Figure 8.**
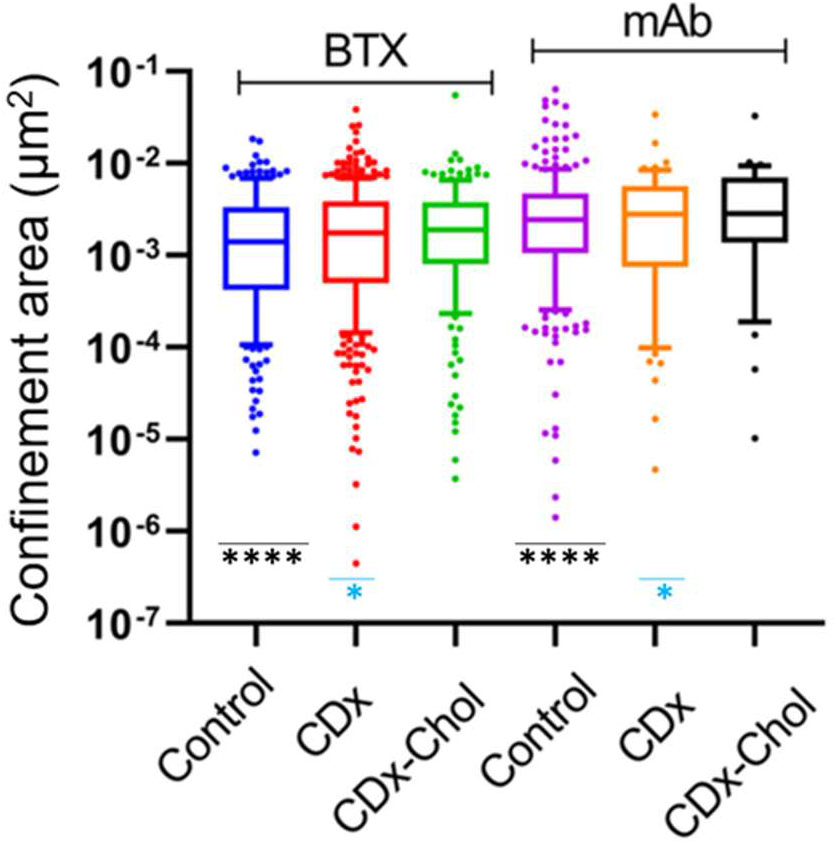
Confinement area (log-scale) for the portions of the trajectories in the obstructed diffusion state. The whiskers represent the interquartile range; the center line is the median; the extremes indicate the 5^th^ and 95^th^ percentile; the dots are outliers. *, p<0.05; ****, <0.0001.

### Diffusion coefficient in the Brownian diffusion state of the two-state model

The diffusion coefficient could be further obtained from application of the corresponding Diffusion Coefficient network (Suppl. Figure 5). The training process was done with Brownian trajectories covering the range of diffusion coefficients between 0.05 and 0.2 μm^2^s^−1^. The network receives a 1-dimensional vector with the normalized squared differences between the adjacent 2-dimensional positions. The network outputs the predicted diffusion coefficient. The architecture is based on a simple convolutional + normalization block, followed by a global max pooling. The pooling is fed to a sequence of two fully connected layers (256 units and 128 units) with a rectified linear unit (ReLU) (Glorot et al., 2010) activation function. For more detailed information see Supplementary Figure 5.

Figure 9 shows the diffusion coefficients of the nAChR under different experimental conditions. They covered a wide range of values, broader and slightly higher for the mAb-labeled receptors than for the BTX-tagged receptors. When we compared the experimental conditions with the same label, differences were observed between BTX control and BTX CDx-Chol (p < 0.05). Peer comparison between BTX and mAb showed very statistically significant differences (p < 0.0001). Similarly, cholesterol reduction (CDx) produced statistically different effects on the diffusion coefficients of BTX- and mAb-labeled receptors (p < 0.001).

**Figure 9.**
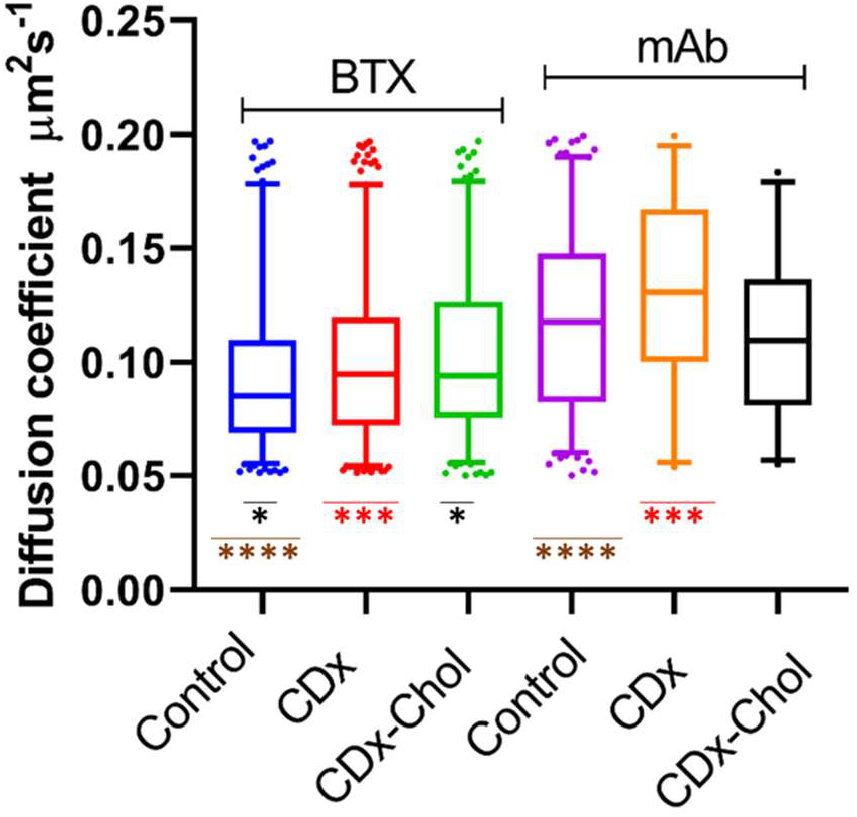
Analysis of the diffusion coefficients of the trajectories classified as two-state by the Classification network (Figure 2). Diffusion coefficients were obtained by application of the Diffusion coefficient Network (Suppl. Figure 5) to the Brownian state segments equal to or greater than 15 steps. The whiskers represent the interquartile range. The center line is the median. The extremes indicate the 5^th^ and 95^th^ percentile, and the dots are outliers. * p < 0.05; ***, p < 0.001; ****, p < 0.0001.

## Discussion

### Heterogeneity of nAChR lateral diffusion

Early static superresolution imaging of the adult muscle-type nAChR using stimulated emission depletion (STED) nanoscopy on fixed cells (Kellner et al., 2007) constituted the first study of a neurotransmitter receptor imaged beyond the diffraction limit in a native membrane environment. The study disclosed the non-random distribution of nAChR nanoclusters at the plasma membrane, imaged at a resolution of ∼ 70 nm.

Our first attempts to follow the dynamics of the receptor protein in live cells made use of single-particle tracking (SPT) and total internal reflection fluorescence (TIRF) microscopy techniques (Almarza et al., 2014). Previous studies from our laboratory using fluorescence recovery after photobleaching (FRAP) and fluorescence correlation spectroscopy (FCS) in the confocal model (Baier et al., 2010) had provided a relatively simple picture of the receptor’s translational diffusion, with the observation of at most two diffusion coefficients. In a previous study from our laboratory (Almarza et al., 2014), it became apparent that receptor mobility was heterogeneous and encompassed a much wider range of diffusional regimes. This could be further dissected using dSTORM-SPT to track the motion of individual nAChR molecules labeled with fluorescent α-bungarotoxin (Mosqueira et al., 2018). Essentially all nAChR single-molecule trajectories exhibited a combination of free Brownian walks interrupted by confinement episodes with average durations of ∼ 400 ms. The STORM study confirmed the heterogeneity of diffusional regimes, from the predominant subdiffusive to the infrequent superdiffusive motion. STORM nanoscopy was subsequently applied to characterize the diffusion of the receptor tagged with the monoclonal antibody mAb35 (Mosqueira et al., 2020). The motional behavior of the nAChRs changed, albeit subtly, suggesting that multivalent antibody-induced crosslinking of the receptor contributes to temporarily hinder its Brownian displacement and confines it in nanoscale domains of ∼34 nm radius. Clustering and immobilization for longer periods into larger domains of ∼ 120-180 nm radii was also observed upon antibody crosslinking. Cholesterol depletion increased the area distended by these transient stop-overs, while cholesterol enrichment had the opposite effect.

### The deep learning multi-network approach

Machine Learning, a broad family of deep learning approaches, has been applied in the last years to characterize single-molecule trajectories. For example, Manzo and coworkers introduced a method based on Random Forest classification algorithms (Breiman, 2001) to sort molecular trajectories according to the most probable theoretical model and to extract the anomalous exponent (Muñoz-Gil et al., 2020). Recently, Shechtman and coworkers (Granik et al., 2019) introduced a set of deep learning methods based on temporal convolutional networks (TCN) (Bai et al., 2018) to characterize SMLM trajectories between 3 diffusion models, namely Brownian, fBm (Sebastian, 1995), and CTRW (Blumen et al., 1984;Chechkin et al., 2009). The TCN architecture consists of two principles: first, the network produces an output of the same length as the input: *ŷ*_0_, …, *ŷ*_*T*_ = *f*(*x*_0_, …, *x*_*T*_) by adding zero padding with a length equal to the kernel size - 1 to maintain the output size. Second, there is no leakage from the future to the past. This is accomplished using causal convolutions, where an output at time *t* can only be affected by elements of time *t*, or the past. In addition, a dilation factor is used in the convolutions to increase the receptive field, allowing the network to have a large history. The overall performance of this type of model to correctly classify the trajectories and estimate parameters such as the diffusion coefficient for short trajectories offers the possibility of extending its application to similar problems. Specifically, the addition of a switching diffusion model (Grebenkov, 2019) combined with obstructed diffusion (Sadegh et al., 2017) and the ability to obtain the segments of each state in individual trajectories has enabled us to produce a more detailed analysis of the transient states during the particle’s diffusion.

To further understand the physical principles underlying the diffusional properties of the nAChR, we implemented Monte Carlo simulations of various physical models of membrane protein diffusion in the plane of the membrane, employing machine learning approaches to simulate a two-state model (Grebenkov, 2019) that combines random Brownian walks with anomalous subdiffusion due to obstacles (Saxton, 1994). We integrated this physical description with other contending models into a multi-network analytical system (Suppl. Figure 1) firstly trained with simulated data and subsequently fed with our experimental data. Initially a set of two classification neural networks was modified and optimized based on the architecture introduced by Shechtman and coworkers (Granik et al., 2019) to specify the most probable model for each trajectory and, when appropriate, the diffusion regime for fractional Brownian motion. We extended the aforementioned architecture to include trajectory segmentation, after a series of modifications to the original network design, allowing us to obtain the state at each step of the trajectory. Having the ability to characterize the segments in each state, we could calculate the diffusion coefficient of each Brownian segment (n > 15) with the aid of a specific neural network that optimized the performance and training stability of the original design (Granik et al., 2019).

A further addition to the multi-network approach presented in this work is the characterization of the diffusion modalities encompassed within the fBm trajectories using the Hurst exponent. The Hurst exponent can be estimated with a traditional approach, e.g. by fitting the mean squared displacement to recover the anomalous exponent. We adopted the methodology of Volpe and coworkers (Bo et al., 2019), selecting the LSTM architecture (Hochreiter and Schmidhuber, 1997). This approach proved to be very convenient in terms of performance and training time. Two additional advantages of this choice were the observed reduction of the estimation error as shown in Suppl. Figure 10, and the ability to determine the overall diffusion range that better characterizes the dynamics of the receptor, namely, subdiffusive, Brownian, or superdiffusive.

To analyze our experimental data we trained more than 3,000 models, each for a specific trajectory length and range of duration. Furthermore, we analyzed the hyperparameters of all the neural networks to increase the accuracy and/or reduce the estimation error (depending on the neural network), and for the short trajectories (n < 50 steps) where the localization error is most apparent.

### The fractional Brownian motion

fBm is a prototypical subdiffusive mechanism (Krapf, 2015) with a Gaussian propagator as in conventional Brownian motion, but with a time-dependent diffusion coefficient (Sebastian, 1995), and with increments that depend on previous increments (Mandelbrot and Van Ness, 1968). To classify the different ranges of fBm diffusion we modified and optimized the original architecture of Shechtman and coworkers (Granik et al., 2019). To minimize the impact of the localization error on short trajectories (∼25-50 steps) the number of convolutional blocks was extended using an intermediate kernel size, allowing us to maintain high precision in the subclassification of the fBm cases. The output of the fBm Subclassification network (Figure 5) indicated that subdiffusive motion with a Hurst index between 0.1 and 0.42 accounted for the behavior of most of the trajectories recorded under control, CDx, or CDx-Chol conditions. The values resulting from the Hurst exponent network (Suppl. Figure 2) did not exhibit significant differences between different experimental conditions, indicating that cholesterol modification did not affect the anomalous exponent that describes subdiffusive trajectories categorized as belonging to the fBm motional regime.

### The two-state model

Those trajectories identified as belonging to the two-state model were fed to the Segmentation network (Figure 7). This network returns the state vector for each trajectory, which provides a flexible solution to extract information about the segments. In previous studies from our laboratory (Mosqueira et al., 2018;Mosqueira et al., 2020) this information was extracted using recurrent analysis and other analytical tools introduced by Krapf and his group to study molecular trajectories (Weigel et al., 2011).

A hypoexponential distribution fitted well the residence times in both states under all experimental conditions studied. The goodness of the fit (> 0.99) with a double-exponential function in essentially all cases (except for the obstructed-diffusion state in the mAb control condition, ∼0.84) suggests that two different processes dictate the residence time in each state, one having a very short duration (in the order of a few ms) and the other with an order of magnitude longer lifetimes. Further experiments will be needed to ascertain the nature of these processes and the underlying biological mechanisms.

The confinement areas corresponding to the segments in the obstructed-diffusion state did not show differences under different experimental conditions for a given probe (BTX or mAb). However, statistically significant differences were apparent when mAb and BTX samples were compared with each other, the latter having smaller confinement areas.

The Brownian segments detected with the Segmentation network (Figure 7), further analyzed with the Diffusion coefficient network (Suppl. Figure 5), showed in all cases higher diffusion coefficients for mAb-tagged samples as compared to BTX-labeled samples and an increase in this metric when cholesterol was reduced under both labeling conditions.

## Conclusions

We developed a deep learning-based multi-network approach to analyze the dynamics of the nicotinic acetylcholine receptor and implemented a two-state model which combines Brownian and obstructed diffusion mechanisms. This implied the addition of a new architecture to segment the two-state trajectories for multiple analysis. To execute each neural network a hyperparameter analysis was performed to obtain a correct generalization for different number of steps and time duration, especially for short trajectories where the localization error greatly affects the accuracy of the models. We next used these models to evaluate the experimental data. From this analysis the conclusion was reached that a two-state model represents more faithfully the great majority of the nAChR trajectories under the present experimental condition.

While we were completing the present work, Manzo and coworkers introduced a Deep Learning approach with a single-hidden layer feedforward network for the anomalous diffusion challenge (www.andi-challenge.org) to tackle the classification, detection of anomalous exponent, and segmentation of trajectories (Manzo, 2021). As part of the “Andy anomalous diffusion challenge” (Muñoz-Gil et al., 2021), a variety of approaches have been presented towards a similar end. The segmentation task only considers one change-point in the trajectory, which may exclude the possibility of multiple transitions in a complex environment, as we show in the present work. Another tactic was recently introduced by Krapf and coworkers using a hidden Markov model to obtain the most probable state at each step for a given trajectory, specifically for a system following a two-state fBm behavior (Janczura et al., 2021). However, given the non-triviality of such task, the problem of trajectory segmentation remains, particularly for short trajectories. Here we present a different approach using Deep Learning for the detection of two states. Future work will compare the performance of our approach with that of the Andy challenge. We can already state that unlike other approaches to tackling the segmentation problem (Muñoz-Gil et al., 2021) our network system is capable of detecting multiple segments of a single molecular trajectory, with one or more change-points.

## Supporting information

Suppl. Mat. Buena Maizon-Barrantes Deep Learning

## Acknowledgments

Thanks are due to Ricardo Di Pasquale for his help in the selection of hyperparameters to generalize the multiple model approach and to Alejo Mosqueira for suggestions regarding the implementation of the two-state model. We acknowledge the support of NVIDIA Corporation for the donation of a TITAN V GPU that facilitated the machine-demanding simulations. Experimental work was supported by grant PICT 2015-2654 IB from the Ministry of Science & Technology of Argentina to F.J.B.

