## Supplementary material for "A deep learning-based approach to model anomalous diffusion of membrane proteins: The case of the nicotinic acetylcholine receptor": Suppl. Mat. Buena Maizon-Barrantes Deep Learning

### Materials

Mouse monoclonal antibody mAb35 (product M-217) against the extracellular moiety of the nicotinic acetylcholine receptor (nAChR) $\alpha_1$-subunit, methyl-β-cyclodextrin (CDx), catalase, glucose oxidase, β-mercaptoethanol, and polyvinylalcohol (PVA, 25,000 MW, prod. No. 184632) were purchased from Sigma Chem. Co. (St. Louis, MO). Alexa-Fluor$^{555}$ -α-bungarotoxin (BTX) and Texas Red-labeled anti-IgG secondary antibodies were purchased from Invitrogen Argentina.

### Methods

#### Cell culture

Cells of the clonal line CHO-K1/A5 (Roccamo et al. 1999) were grown in Ham's F12 medium supplemented with 10% fetal bovine serum for 2-3 days at 37°C before experiments.

#### Acute cyclodextrin-mediated cholesterol depletion/enrichment of cultured cells

Acute cholesterol depletion or enrichment was performed prior to fluorescent labeling by treating CHO-K1/A5 cells with 10-15 mM CDx or CDx-cholesterol (CDx-Chol) complexes in Medium 1 ("M1": 140 mM NaCl, 1 mM $CaCl_2$, 1 mM $MgCl_2$ and 5 mM KCl in 20 mM HEPES buffer, pH 7.4) (Borroni et al. 2007; Almarza et al. 2014). Samples were taken at 20 min from culture dishes incubated at 37°C in the presence or absence of the cholesterol-modifying chemical.

#### Single-molecule localization superresolution microscope setup

The optical nanoscope constructed in our laboratory was operated in the stochastic optical reconstruction microscopy (STORM) (Rust et al. 2006) modality, as described in detail in ref. (Mosqueira et al. 2018).

#### Cell-surface fluorescence staining of nAChRs

CHO-K1/A5 cells grown on 18 mm diameter No. 1.5 glass coverslips (WRL) in Ham's F12 medium at 37°C were washed thrice with M1 medium and incubated with Alexa-Fluor$^{555}$- BTX for 45 min at 4°C as in ref. (Mosqueira et al. 2018). In the case of antibody labelling, cells were incubated with mAb35 primary antibody under similar conditions, washed thrice with cold M1 containing 10% bovine foetal serum, and incubated with Texas Red-labelled goat anti-mouse secondary antibody for 1 additional h at 4°C in M1 medium as described in (Mosqueira et al. 2020).

#### Single-molecule superresolution imaging

STORM microscopy was carried out as described previously (Mosqueira et al. 2018; Mosqueira et al. 2020) using a planapochromatic TIRF 100x, 1.49 N.A. oil immersion objective in conjunction with a back-illuminated electron multiplying CCD camera (iXon-Plus DU-860, Andor Technology, Belfast, Northern Ireland) set to acquire a stream of images at maximum frame rate at a gain of ~250 and 3.6 photoelectrons per A/D count. Streaming movies were acquired from between 8 and 15 cells using the software SlideBook (Intelligent Imaging Innovations, Boulder, CO) and exported as 16-bit TIF or Matlab files for subsequent off-line analysis. All images were recorded from the ventral, coverslip-contacting membrane of the cells. Cells were inspected after image acquisition to ensure preservation of cell morphology.

### Superresolution data analysis

#### i) Determination of sub-diffraction molecular coordinates

The off-line localization of the x,y coordinates of the nAChR spots was carried out using the image analysis package ThunderSTORM (https://code.google.com/p/thunder-storm/) (Ovesný et al. 2014) run as a plugin in ImageJ (https://imagej.nih.gov/ij/). ThunderSTORM is particularly suitable for separating multiple overlapping PSFs (typical emitter density was 5 localizations per frame). In order to account for the discrete nature of pixels in digital cameras, an integrated-form of a symmetric 2D Gaussian function was fitted to the spots using Levenberg–Marquardt least-squares minimization routines. Localizations that were too close together to be independent were discarded. The ThunderSTORM multi-emitter fitting analysis was enabled, and the limiting intensity range was set at 500-2,000 photons. Other ThunderSTORM filters were enabled to remove uncertainty-based duplicates (e.g. multiple emitters and duplicate localizations, as described in ref. (Huang et al. 2011)). Lateral drift was estimated experimentally using fiducial 100 nm coverslip-adhered fluorescent beads and corrected via the appropriate filter in ThunderSTORM. Localization precision was calculated automatically using a modified version of the formula in ref. (Thompson et al. 2002) which considers the EM gain of the EM-CCD camera (Quan et al. 2010), through the following expression:

$$\langle(\Delta x)^2\rangle = \frac{2\sigma^2 + a^2/12}{N} + \frac{8\pi\sigma^4 b^2}{a^2 N^2} \qquad \text{(Eq. 1)}$$

where $\sigma$ is the standard deviation of the fitted point spread function (PSF), $a$ is the pixel size in nm, $N$ is the intensity expressed in number of photons and $b$ is the background signal level in photons. The average localization precision was 40 nm.

**ii) Single-particle tracking (SPT)**

The fluorescent particles were detected by a generalized likelihood ratio test algorithm specifically designed to detect point spread function-shaped (i.e. Gaussian-like) spots using ThunderSTORM (Ovesný et al. 2014) and exported in a format suitable for tracking analysis using an ad-hoc MATLAB routine written in our laboratory. Detected (validated) particles were further analyzed for their trajectories with the software package Localizer (https://bitbucket.org/pdedecker/localizer) (Dedecker et al. 2012) implemented in Igor Pro (Wavemetrics Inc. https://www.wavemetrics.com). Two critical parameters were set in Localizer: the maximal number of frames (3 = 30 ms and 2= 20 ms for BTX- and mAb-labeled samples, respectively) that a given molecule was allowed to blink ("Max blinking"), and the maximum distance ("Max jump distance") at which two points could lie and be attributed to the same trajectory (3 pixels). The optimal "Max blinking" ($t_{off}$) was determined following the method of Annibale and coworkers (Annibale et al. 2011) which implies that the number of photoblinking fluorescent molecules $N$ in the sample can be estimated from the number of counts (localizations) at different dark times $t_d$, $N(t_d)$, by fitting to the semi-empirical equation:

$$N(t_d) = N\left(1 + n_{blink}e^{\left(\frac{1-t_d}{t_{off}}\right)}\right) \qquad \text{(Eq. 2)}$$

in the regime of low dark time $t_d$ values (Annibale et al. 2011). The maximum distance for the merging filter in our experiments was determined from the camera pixel size and the distance-filter criterion described by others (Lu et al. 2014). Briefly, localized molecules that reappeared in consecutive frames were considered as corresponding to the same molecule if the frame-to-frame displacement (tracking radius) was within 106 nm, thus allowing the monitoring of molecules with diffusion coefficients of up to 1.33 $\mu m^2 s^{-1}$, i.e. conservatively higher than the upper bound for nAChR nanocluster diffusion estimated from TIRF-SPT experiments in our laboratory (Almarza et al. 2014). Following the above argument, the "Max jump distance" was deduced by considering the maximum distance that a nAChR is allowed to "jump" within the merge time window established above, i.e. the maximum distance that, on average, a single-molecule can travel in 2 or 3 frames for mAb- and BTX-labeled samples, respectively.

**iii) Exclusion of immobile particles**

Stationary (immobile) molecules were excluded from the analysis of single-molecule trajectories following a series of recently introduced criteria (Golan and Sherman 2017). The procedure sets a threshold value on the ratio of the radius of gyration $\mathrm{R_g}$ and the mean step size $|\Delta \mathrm{r}|$ of the particles´displacement. In the case of ideal immobile particles this ratio is constant, whereas for mobile particles the ratio increases. The normalized ratio:

$$(\sqrt{\pi/2}\ \ (\mathrm{R_g}/\langle|\Delta \mathrm{r}|\rangle) \qquad \text{(Eq. 3)}$$

was obtained from experiments with 4% paraformaldehyde-fixed cells, and the ratio was subsequently employed to obtain the threshold value applied to live cell experiments. Golan and Sherman (2017) discuss the advantages of this method over the use of the diffusion coefficient or $\mathrm{R_g}$ alone for excluding immobile particles; the two latter procedures would falsely classify immobile particles as mobile. Threshold values above the 95$^{th}$ percentile were obtained by pooling data from different cells in independent sets of experiments and a mid-value was selected.

#### Statistical analyses

Statistical data are expressed as the mean ± 95% confidence interval (CI) unless specified otherwise. One-way analysis of variance followed the Kruskal-Wallis test or Sidak's multiple comparisons test implemented with the GraphPad Prism software. The one-sample D'Agostino & Pearson test was applied to assess whether the data were normally distributed or not. To compare two distributions, we use the Kolmogorov-Smirnov test for two samples.

### Supplementary Results

#### Neural Networks training and diffusion models

Networks were trained using simulated data. The process starts by simulating a training set, and a new and randomly generated validation set is used foreach epoch to avoid any bias in the training process.

Trajectories are simulated in two steps; first, the specific model generates a random trajectory, and an adaptation to the experimental pixel size (106 nm) of the actual microscopy experiments is performed when appropriate. A random offset is added next to the x,y positions, and a localization error at each step is randomly generated and added to the coordinates' value. This error is generated using a normal distribution with a mean value of 40 nm and a standard deviation of 10 nm, also based on the localization error of the actual experimental setup (see above). The simulation of all the trajectories is based on independent axes, the simulated axes data being joined together to form the 2D trajectory.

Recently, Shechtman and coworkers (Granik et al. 2019) implemented a deep learning approach based on neural networks to classify single-particle trajectories based on their diffusion behavior: Brownian (free diffusion) and the anomalous diffusion (Höfling and Franosch 2013) models, fractional Brownian motion (fBm) (Feder et al. 1996), and continuous time random walk (CTRW) (Grebenkov and Tupikina 2018; Chechkin et al. 2009). These models represent different modalities of a particle's translational motion in a 2-dimensional space. The explored region as a function of time can be expressed with the generalized formulation of the mean-squared displacement:

$$\langle r^2 \rangle = Kt^{\alpha} \qquad \text{(Eq. 4)}$$

Where $K$ is the generalized diffusion coefficient, and $\alpha$ is the anomalous exponent. The sublinear case $\alpha < 1$ indicates subdiffusion, $\alpha = 1$ is the "normal", or Brownian diffusion, and $\alpha > 1$ is the superdiffusion range.

The idea behind the neural network approach was to achieve greater accuracy and be able to use shorter trajectories (< 25 steps) than those required for time-averaged mean-square displacement (TAMSD) analysis:

$$TAMSD = \frac{1}{N-m}\sum_{i=1}^{N-m}\left[x_j(t_i + m\Delta t) - x_j(t_i)\right]^2 \qquad \text{(Eq. 5)}$$

where $x_j$ is the position sampled at $N$ discrete times $t_i = i\Delta t$, $\Delta t$ is the acquisition time (in our case, $\Delta t = 10ms$ ) and $i$ the frame number.

The first model, fractional Brownian motion (fBm) is a generalization of Brownian motion, where the increments depend on the previous increments and has the following covariance function:

$$Cov(W_t, W_s) = \frac{1}{2}(|t|^{\alpha} + |s|^{\alpha} - |t-s|^{\alpha}), \quad t, s \geq 0 \qquad \text{(Eq. 6)}$$

The exponent $\alpha$ is usually replaced with the Hurst exponent: $H = \frac{\alpha}{2};\ \ H \epsilon\ (0,1)$. This index indicates the diffusion regime, where $H < 0.5$ is observed in subdiffusive motion, $H = 0.5$ implies Brownian motion, and $H > 0.5$ corresponds to superdiffusion. In the simulation of this model, and also for the continuous time random walk (CTRW) model (see below), we used the algorithms of Shechtman and coworkers (Granik et al. 2019). In the case of fBm the circulant embedding approach (Kroese and Botev 2015) was used. This algorithm requires the Hurst exponent to be defined before the simulation, allowing us to use a supervised-learning approach to train the neural nets.

CTRW describes a process consisting of a series of independent, identically distributed random jumps separated by waiting times. These jumps were simulated using the Symmetric Lévy α-stable probability density function (PDF):

$$\xi_{\alpha} = \gamma_x \left(\frac{-ln\, u\, cos\phi}{cos((1-\alpha)\phi)}\right)^{1-1/\alpha} \frac{sin(\alpha\phi)}{cos\phi} \qquad \text{(Eq. 7)}$$

Where $\phi = \pi(v - 1/2)$, and $u, v\ \epsilon\ (0,1)$. For the waiting times, we employed the Mittag-Leffler PDF:

$$\tau_{\beta} = -\gamma_t ln\, u\, \left(\frac{\sin(\beta\pi)}{\tan(\beta\pi v)} - \cos(\beta\pi)\right)^{\frac{1}{\beta}},\ u, v\ \epsilon\ (0,1) \qquad \text{(Eq. 8)}$$

In both cases, $u$ and $v$ are generated randomly, with $\gamma$ fixed to 1 and $\beta$ to 0.5.
In this work we introduce a third model which combines switching diffusion (Weron et al. 2017; Grebenkov 2019) and obstructed diffusion (Sadegh et al. 2017; Weigel et al. 2012) modalities of translational diffusion. Switching diffusion consists of a set of N multiple states, each one with a defined diffusion coefficient $D_i$, and a set of switching rates $k_{ij}$ that represent the rate of the random transitions from a state $i$ to a subsequent state $j$ (Weron et al. 2017; Grebenkov 2019). These states may represent different conformations of a macromolecule, or a temporal binding to other molecules, resulting in a heterogeneous process.

**Software packages**

All the scripts, simulations, and neural networks were developed using Python and some of its packages like Numpy, Keras and SciPy, among others. To analyze and plot the results Matplotlib, MATLAB, and GraphPad Prism were used.

**Two-state diffusion simulation**

The model is based on a more general two-state switching diffusion simulation introduced by Grebenkov for Brownian trajectories alternating between two states (Grebenkov 2019). We adapt this general scheme by introducing an obstructed diffusion state (Sadegh et al. 2017; Weigel et al. 2012) to model in silico the possible contribution of the latter to nAChR dynamics at the cell surface. The diffusing particle alternates between 2 states, a Brownian diffusion state with a diffusion coefficient *D*, and an obstructed diffusion state. A characteristic residence time is defined within an exponential distribution random generator to generate the mean residence time for each state in the switching diffusion (Grebenkov 2019). To cover a wide range of values of residence times (Mosqueira et al. 2018), before running each simulation the mean value for the generator is also selected randomly. The mean residence time is expressed in terms of the switching rate $\mathrm{t_i} = \frac{1}{k_i}$.
Table 1 below shows the range of switching rates used to randomly generate the mean residence time of the sojourns in the obstructed state, expressed in $\frac{1}{frames}$.

| Switching rate | Low | High |
|---|---|---|
| $k_0$ (Brownian diffusion state) | 0.01 | 0.08 |
| $k_1$ (Obstructed diffusion state) | 0.007 | 0.2 |

Supplementary Table 1. Rate constants for the calculation of the mean residence time of the confined sojourns.

With the generated residence times, the simulation starts by choosing a random initial state. In the case of Brownian diffusion, the diffusion coefficient *D* is randomly chosen from the range [0.05-0.2], expressed in $\mu m^2 s^{-1}$. *D* is kept constant over the entire simulation of the trajectory. The steps in the Brownian motion state are randomly generated Gaussian increments with μ=0 and σ=1, re-scaled by $\sqrt{2D\Delta t}$, where Δt is the duration in seconds divided by the number of steps of the trajectory. Then, the increments are cumulatively summed up.

In the case of the obstructed diffusion state, the simulation commences by generating a random squared fence with a side length in the range [20-40] nm. In this case, the step sizes are drawn from a normal distribution with standard deviation σ=5. Permeation of the obstacles impeding the particle's diffusion occurs when the residence time ends. In those cases where a new step is outside the fence, the particle remains in the old positions until a new valid step occurs.

**Neural networks architecture and development**

Our approach draws on the architecture introduced by Shechtman and coworkers (Granik et al. 2019) based on Temporal Convolutional Networks (TCN) (Bai et al. 2018) and we expanded the idea to new applications such as trajectory segmentation. In other cases, an architecture based on Long-Short Term Memory networks (Bo et al. 2019) was used to predict the Hurst Index for trajectories classified as fractional Brownian model (fBm). To obtain the diffusion coefficient *D*, we combined concepts of TCN (Bai et al. 2018) and introduced data transformation to the neural network input.  This increased the robustness of the training process for the particular case of our-simulated data.

All the networks were optimized for the range of steps of our experimental data. To accomplish this task, a hyperparameter optimization was performed, using a Grid Search algorithm (Yang and Shami 2020). This analysis yielded good results even with short trajectories where the localization error greatly affects the network performance. The dataset was distributed with an 80:20 ratio for training and validation data, using approximately 50.000 to 60.000 trajectories. The hyperparameter study, and the more than 3000 models trained to evaluate our data required more than 20 days non-stop NVIDIA TITAN V GPU time. This included testing approximately 70 different setups for each neural network to select the correct value for each parameter of the neural network optimization algorithm.

The input for all the implemented neural networks, except for the Diffusion Coefficient Network (Suppl. Figure 5), is a 1D tensor with an individual axis. The network output for a 2D trajectory is the mean value of the network's output tensor of the individual coordinates. This analysis considers each trajectory's dimension independently from the others.

The multi-network pipeline in Suppl. Figure 1 separates our data according to the previous network results. In the case of the Hurst Exponent Network, we use the fBm diffusion range obtained using the fBm classification network to select a specific network trained for the detected range, thus reducing the prediction error compared to a complete range network [0,1].

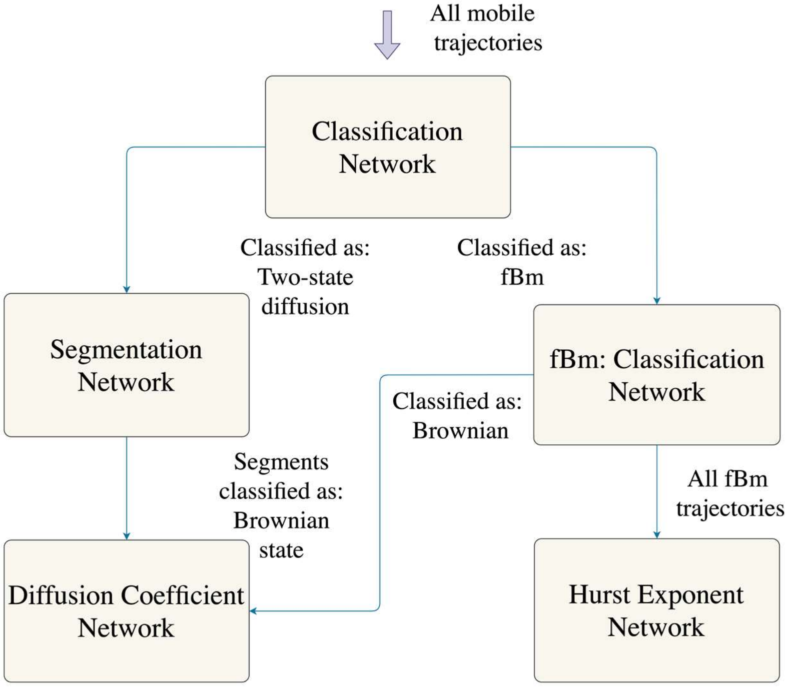


Supplementary Figure 1. General outline of the classification networks designed and applied in this work to test three diffusion physical models. The input is designated "all mobile trajectories". These trajectories result from application of the criteria of Golan and Sherman (Golan and Sherman 2017), which exclude those trajectories in the set of raw validated trajectories that are considered immobile. The first classification network selects the most probable model. The fBm classification network only categorizes those trajectories that fall within the range set for fBm. The output data are used for the selection of a specialized Hurst exponent network within the detected range. In the case of the two-state diffusion model, the segmentation network selects the most probable state for each step in the trajectory and the output data are used to compute the confinement area and residence time of the sequences in this state. The Brownian state sequences are used as input for the Diffusion Coefficient Network where the value of each individual sequence is predicted.

The first network is the Physical Model classification network (Figure 2). Its inputs are the trajectories, and it classifies the data among the three models considered, namely fBm, CTRW, and two-state diffusion. The output assigns a probability to each category, and the one with the highest value is chosen. This result indicates which path the trajectory follows in the pipeline. Supplementary Table 2 shows the results of applying the Physical Model classification network of Figure 2 to the complete dataset of mobile nAChR trajectories (25 to 900 steps).

Table 2. Output data of the Physical Model classification network (Suppl. Figure 1 and Figure 2 in main text)

| | fBm | CTRW | two-state |
|---|---|---|---|
| | BTX | | |
| **Control** | 17.9 (13.5 - 38.0) | 2.9 (2.2 - 26.7) | 79.2 (59.8 - 84.3) |
| **CDx** | 21.9 (16.5 - 41.3) | 5.6 (4.2 - 29.1) | 72.5 (54.5 - 79.3) |
| **CDx-Chol** | 9.4 (7.0 - 32.3) | 1.9 (1.4 - 26.7) | 88.7 (66.3 - 91.6) |
| | mAb | | |
| **Control** | 8.0 (6.0 - 31.4) | 3.2 (2.4 - 27.9) | 88.8 (66.1 - 91.6) |
| **CDx** | 6.9 (5.2 - 30.5) | 2.8 (2.1 - 27.3) | 90.3 (67.5 - 92.7) |
| **CDx-Chol** | 19.6 (14.8 - 39.3) | 6.9 (5.2 - 29.7) | 73.5 (55.5 - 80.0) |

If a trajectory is classified as fractional Brownian motion, the fBm classification network (Figure 4) selects next the most probable type of diffusion between subdiffusive, Brownian, and superdiffusive. The output also assigns a probability to each diffusion type, and the highest is selected. The architecture shares similarities with the classification network (Figure 2) but differs from the latter in that a new set of convolutional blocks was added to increase the performance, and the number of units in the dense layers was scaled. The range for the different diffusion modes was selected using the following values for the Hurst Index: subdiffusive [0.1-0.42], Brownian [0.43-0.57], and superdiffusive [0.58-0.9]. Supplementary Table 3 shows the output of the fBm classification network in Figure 4. The results indicate most of the trajectories are categorized as subdiffusive, with a Hurst exponent lower than 0.42, for both BTX- and mAb-labelled samples.

Supplementary Table 3. Output of the fBm classification network (Figure 4). Input data are the trajectories classified as fBm by the Physical Model network (Figure 2). The results are expressed as percentages.

| | fBm subdiffusive | fBm Brownian | fBm superdiffusive |
|---|---|---|---|
| | BTX | | |
| **Control** | 86.3 (72.0 - 88.5) | 11.3 (9.4 - 25.9) | 2.5 (2.1 - 18.6) |
| **CDx** | 77.9 (65.4 - 81.4) | 20.8 (17.5 - 33.4) | 1.3 (1.1 - 17.1) |
| **CDx-Chol** | 75.0 (63.4 - 78.9) | 15.0 (12.7 - 28.1) | 10.0 (8.5 - 23.9) |
| | mAb | | |
| **Control** | 75.7 (64.0 - 79.4) | 18.9 (16.0 - 31.5) | 5.4 (4.6 - 20.0) |
| **CDx** | 100.0 (83.5 - 100.0) | 0.0 (0.0 - 16.5) | 0.0 (0.0 - 16.5) |
| **CDx-Chol** | 75.0 (63.5 - 78.8) | 25.0 (21.2 - 36.5) | 0.0 (0.0 - 15.3) |

The outcome of the fBm classification network is used as a priori information to select the range for the Hurst Exponent network (Suppl. Figure 2), allowing for more specific training. The Hurst Exponent network has a different architecture from the rest. We used the structural design introduced by Bo and coworkers (Bo et al. 2019) with the aforementioned modification of the Hurst range values. The network output is the predicted value of the Hurst index. The architecture comprises two Long-Short Term Memory (LSTM) (Hochreiter and Schmidhuber 1997) layers, followed by a fully connected Scaled Exponential Linear Unit (SELU) (Klambauer et al., 2017) activation function and the output layer.

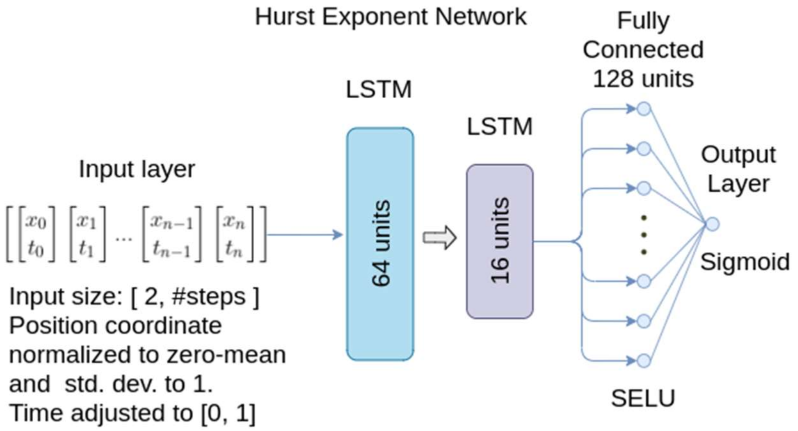


Supplementary Figure 2. Hurst Exponent Network based on the architecture described in ref. (Bo et al. 2019). The network output is optimized for a specific fBm diffusion range -subdiffusive, Brownian, and superdiffusive- allowing reduction of the prediction error with respect to an output for the complete range network [0-1]. This is possible because of the prior information provided by the network in Figure 4, which determines the diffusion range.

Supplementary Table 4. Mean and 95% confidence interval of the mean after applying the Hurst Exponent Network over the entire range (25-900 steps) of fBm trajectories.

| **Hurst Exponent Subdiffusive trajectories** | **Control** | **CDx** | **CDx-Chol** |
|---|---|---|---|
| **BTX** | 0.296 ± 0.007 | 0.296 ± 0.006 | 0.291 ± 0.012 |
| **mAb** | 0.279 ± 0.015 | 0.281 ± 0.018 | 0.289 ± 0.021 |

The trajectories classified as two-state by the initial Classification Network (Figure 2) were further analyzed using the Segmentation Network (Figure 7). This network was trained using two-state diffusion simulated data. Only cases with 1 or more switching of states were considered. This condition is also observed in the simulated data because in some cases a simulated residence time is equal to or longer than the duration of the simulated trajectory, resulting in a fully Brownian or obstructed diffusion motion without a percolation motion.
The segmentation network (Figure 7) outputs a 1D tensor with a value between 0 and 1 for each step. A value < 0.5 indicates a Brownian state, > 0.5, an obstructed diffusion state. Segments sharing the same individual state of each step are detected and stored as data for future computation of residence times and confinement area.
The residence times were computed using the data from the frame numbers, the frame rate, and the segments obtained from the state tensor. The distribution of the residence times is shown in Supplementary Figures 3 and 4. A plot of the fitted hypoexponential distribution was added in each case, the PDF following the equation:

$$PDF(x) = \frac{\lambda_2}{\lambda_2 - \lambda_1}\, \lambda_1\, e^{-\lambda_1 x} + \frac{\lambda_1}{\lambda_1 - \lambda_2}\, \lambda_2\, e^{-\lambda_2 x} \qquad \text{(Eq. 9)}$$

In both cases, $\lambda_i$ rates are positive values and $\lambda_1 \neq \lambda_2$.
The confinement area for the obstructed diffusion state is obtained as the difference between the maximum and minimum coordinate values of the segment.

Supplementary Table 5. Median and interquartile range of the confinement areas in the obstructed diffusion state. The lower value between parentheses is the upper bound of the first quartile and the other value is the lower bound of the fourth quartile.

| **Confinement area ($\mu m^2$)** | **Control** | **CDx** | **CDx-Chol** |
|---|---|---|---|
| **BTX** | 0.0014 (0.0004 - 0.003) | 0.0017 (0.0005 - 0.004) | 0.0019 (0.0008 - 0.004) |
| **mAb** | 0.0024 (0.0011 - 0.005) | 0.0028 (0.0007 - 0.006) | 0.0028 (0.0014 - 0.007) |

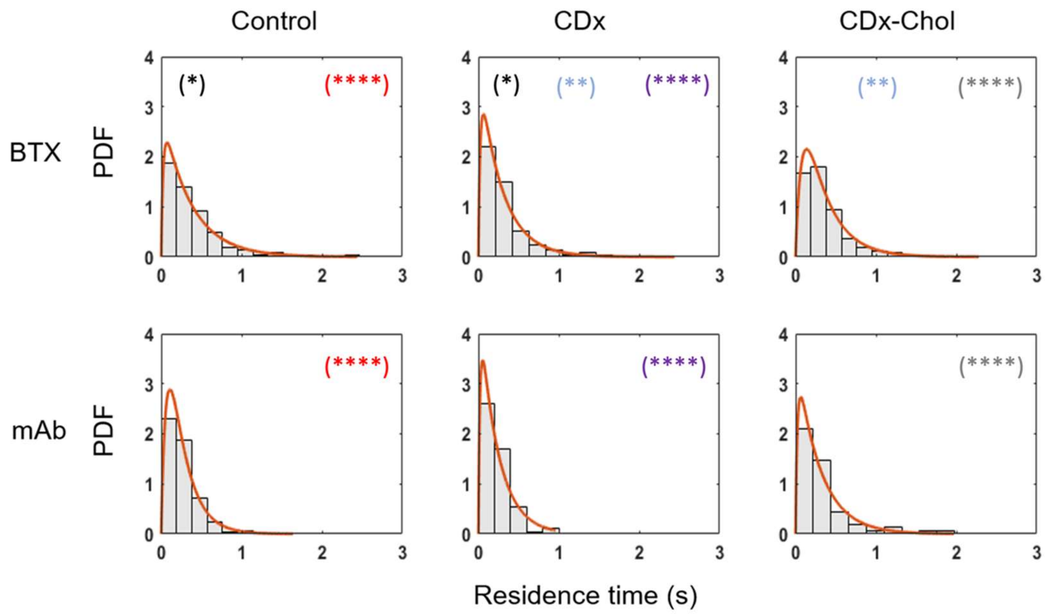


Supplementary Figure 3. Residence times for the trajectories that undergo Brownian motion. The brown line is the curve corresponding to a hypoexponential fit. *, $p < 0.05$; **: $p < 0.01$; ****, $p < 0.0001$.

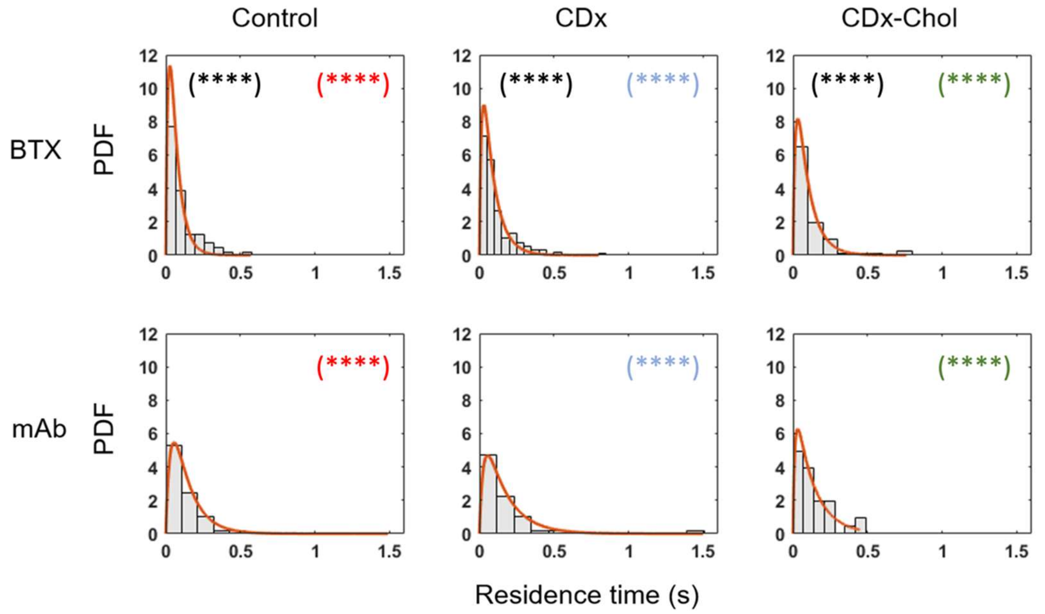


Supplementary Figure 4. Residence times for the sequences in the obstructed diffusion state. The brown line is the curve corresponding to a hypoexponential fit. **** is $p < 0.0001$.

Finally, the Diffusion Coefficient Network (Suppl. Figure 5) receives a Brownian trajectory or sequence from a two-state model and predicts the value of *D*. The network was trained with trajectories in the range [0.05- 0.2] $\mu m^2 s^{-1}$, which defines the boundaries it can detect. To train this network, Brownian trajectories were simulated using the same algorithm used for the Brownian state in the two-state model.

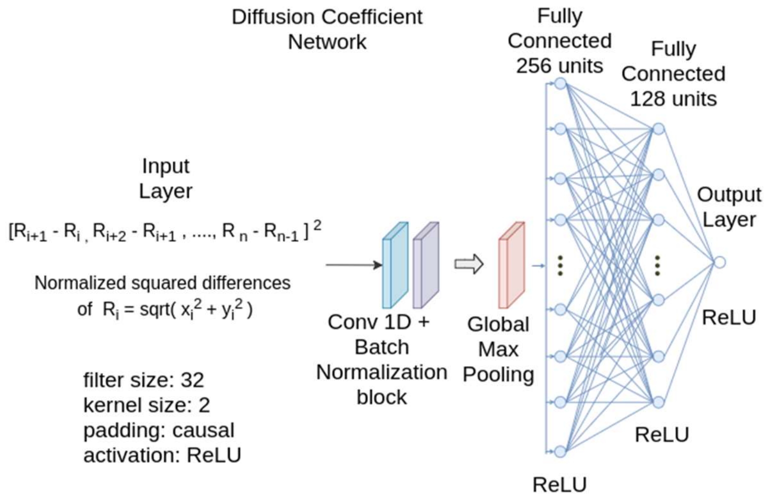


Supplementary Figure 5. Diffusion Coefficient network. This network receives the normalized squared differences of each 2D coordinate and outputs the diffusion coefficient. The architecture is similar to that of ref. (Granik et al. 2019) but with changes in the input and output to increase the performance in the range of steps and localization error characteristic of our data. A specific range of values [0.05-0.2 $\mu m^2 s^{-1}$] of *D* was selected to train the network.

Supplementary Table 6. Median and the 95% CI lower and upper limit for the median of the diffusion coefficients calculated for the Brownian portions of the trajectories by the Diffusion Coefficient network (Suppl. Figure 5). The network was trained to detect trajectories in the range 0.05 to 0.2 $\mu m^2 s^{-1}$.

| **Two-state model - Brownian state** | |
|---|---|
| **Diffusion coefficient [$\mu m^2 s^{-1}$]** | |
| **BTX** | |
| Control | 0.085 (0.079 – 0.092) |
| CDx | 0.095 (0.089 – 0.098) |
| CDx-Chol | 0.094 (0.09 – 0.103) |
| **mAb** | |
| Control | 0.118 (0.11 – 0.126) |
| CDx | 0.131 (0.104 – 0.153) |
| CDx-Chol | 0.109 (0.086 – 0.132) |

**Metrics of the transitions in the two-state model**

The switching behavior of the two-state model raises the question as to the number of transitions between the states. A trajectory could start with a Brownian diffusion and, after any number of steps, switch to obstructed diffusion or vice versa. A difference in the distribution of the number of transitions could indicate a bias of the model in the prediction of the state, i.e. detect only one type of transition. Also, a higher number of transitions might indicate a high number of confinement regions.

As shown in Suppl. Figure 6, in the case of the trajectories of mAb-labeled receptors, the maximum number of transitions was 2 whereas for BTX-labeled nAChRs up to 4 transitions could be observed. Comparison of the experimental conditions of samples labeled with the same probe, and between BTX and mAb, did not show statistically significant differences.

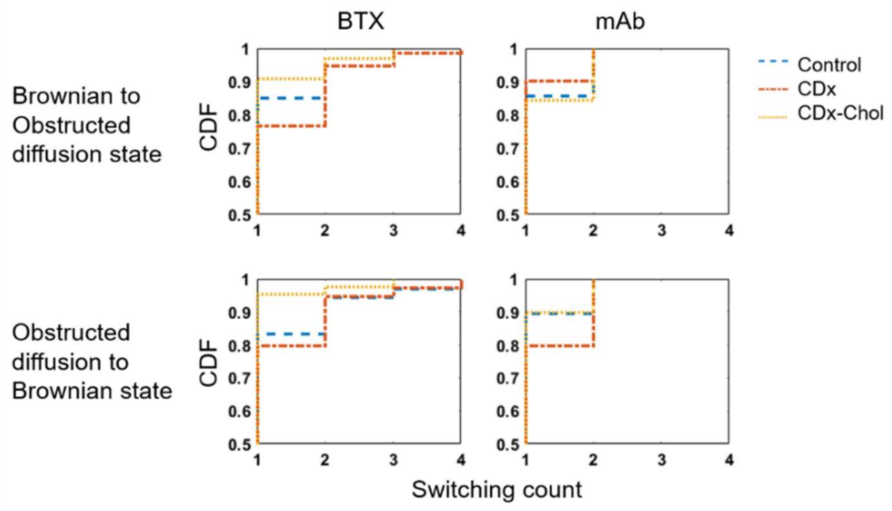


Suppl. Figure 6. Empirical cumulative distribution function of the switching count for transitions between states in the two-state model of Figure 7 in the main text.

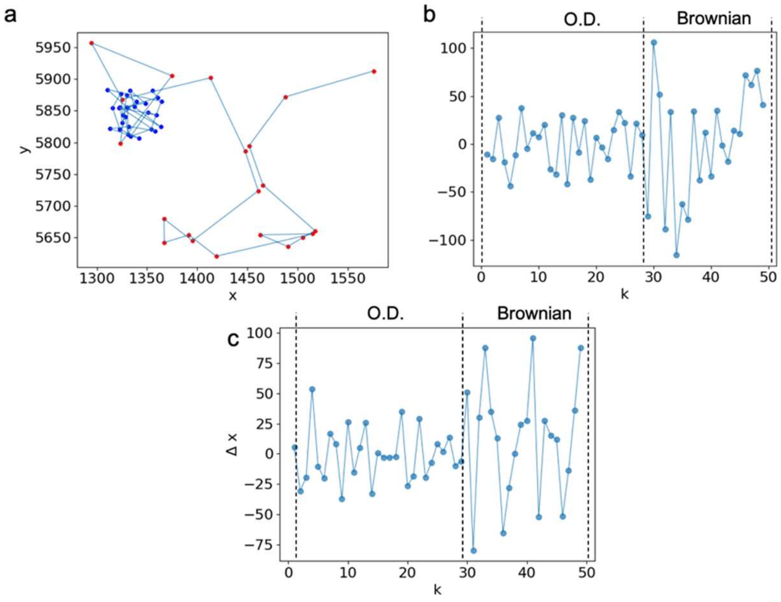


Suppl. Figure 7. a) Predicted state for each step of the trajectory after evaluation using the Segmentation network (Figure 7). Obstructed diffusion is shown in blue, and the red dots correspond to Brownian diffusion; b-c) the variation between positions $k = p_{i+1} - p_i$, for each coordinate. A clear visual difference can be observed between the obstructed diffusion state and the Brownian state.

**Neural networks optimization and accuracy**

A hyperparameter optimization was performed to define a setup able to correctly generalize particularly in the range of 25 to 50 steps, where the localization error significantly affects the accuracy of the neural networks. Briefly, the study starts by defining a list of possible values for each parameter of the Adaptive Moment Estimation (Adam), the stochastic optimization algorithm (Kingma and Ba 2014). Next, a Grid Search (Yang and Shami 2020) is performed, and the validation loss is plotted for all the possible solutions. The space of solutions is reduced by selecting a range where the lowest losses were found, culminating in the selection of the best 10 setups. This procedure is repeated twice, for 25 and 50 steps-long trajectories. The final setup is selected by comparing the two lists and choosing the option that appears best for both lengths of trajectories. The purpose of this procedure is to define a correct generalization of the hyperparameters.

During the training process, the validation set is composed of 50% constant trajectories over all possible solutions and the remaining 50% is a randomly generated epoch. This approach was selected to avoid a bias in the selection of the correct setup.

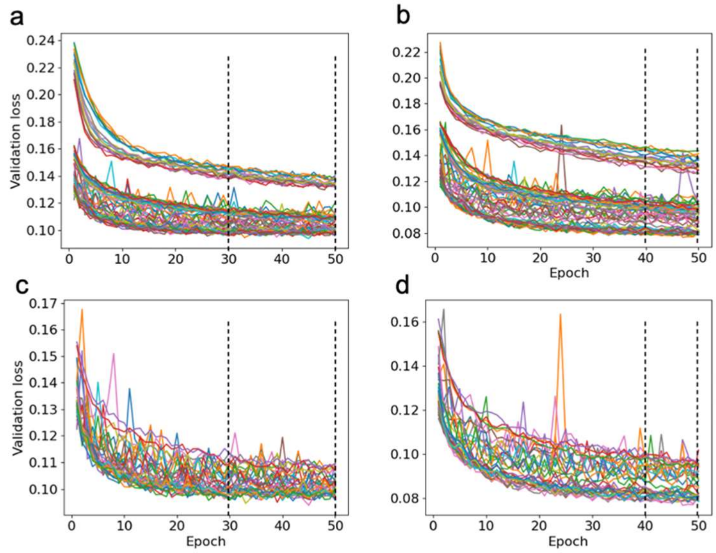


e

| | Learning Rate | Batch size | AMS grad | Epsilon | Mean Loss | S.D. |
|---|---|---|---|---|---|---|
| 25 steps | 0.001 | 16 | FALSE | 1,00E-06 | 0.098 | 0.002 |
| | 0.001 | 16 | FALSE | 1,00E-07 | 0.098 | 0.001 |
| | 0.001 | 16 | FALSE | 1,00E-08 | 0.098 | 0.001 |
| | 0.001 | 32 | FALSE | 1,00E-07 | 0.098 | 0.001 |
| | 0.001 | 8 | TRUE | 1,00E-08 | 0.098 | 0.001 |
| | 0.001 | 16 | TRUE | 1,00E-06 | 0.098 | 0.001 |
| | 0.001 | 16 | TRUE | 1,00E-08 | 0.098 | 0.002 |
| | 0.001 | 32 | TRUE | 1,00E-06 | 0.098 | 0.002 |
| | 0.001 | 32 | TRUE | 1,00E-07 | 0.099 | 0.001 |
| | 0.001 | 32 | TRUE | 1,00E-08 | 0.099 | 0.001 |
| 50 steps | 0.001 | 16 | FALSE | 1,00E-06 | 0.079 | 0.002 |
| | 0.001 | 16 | FALSE | 1,00E-07 | 0.079 | 0.001 |
| | 0.001 | 16 | FALSE | 1,00E-08 | 0.079 | 0.001 |
| | 0.001 | 32 | FALSE | 1,00E-07 | 0.080 | 0.001 |
| | 0.001 | 16 | TRUE | 1,00E-06 | 0.080 | 0.001 |
| | 0.001 | 16 | TRUE | 1,00E-07 | 0.080 | 0.002 |
| | 0.001 | 16 | TRUE | 1,00E-08 | 0.081 | 0.002 |
| | 0.001 | 32 | TRUE | 1,00E-06 | 0.081 | 0.001 |
| | 0.001 | 32 | TRUE | 1,00E-07 | 0.081 | 0.001 |
| | 0.001 | 32 | TRUE | 1,00E-08 | 0.081 | 0.001 |

Suppl. Figure 8. Example of the hyperparameter optimization procedure. a-b) show all possible combinations of hyperparameters explored. The dashed black lines indicate the first selected range to reduce the solutions' space; c-d) represent the reduction to 50% of the best validation loss solution´s space; also in this case the dashed lines represent the second selected range; e) the table contains the final 10 best results for 25 and 50 steps, respectively. The orange-colored rows represent the set of hyperparameters that best generalize between 25 and 50 steps. These sets were used to train all the models of the network.

The confusion matrices in Suppl. Figure 9 show the accuracy of the classification neural network (Figures 2 and 4) and the segmentation network (Figure 7) used to correctly detect the class/state for each model.

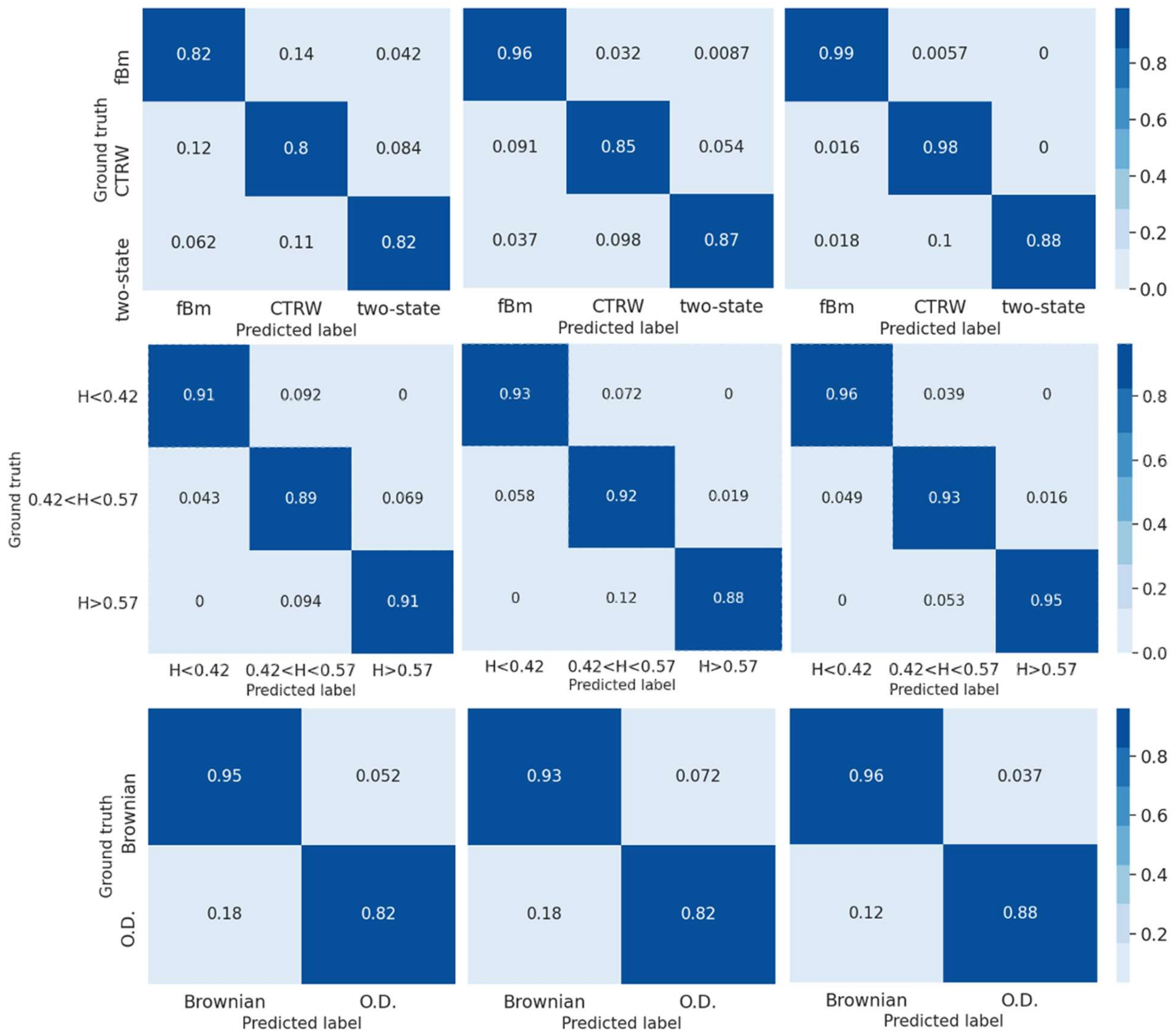


Suppl. Figure 9. Confusion matrices of a random validation set of 10.000 trajectories analyzed with the Classification Network (Figure 2) (upper row), the fBm Subclassification Network (Figure 4) (center row), and the Segmentation Network (Figure 7) (lower row). From left to right, each matrix represents the results obtained for trajectories of 25, 50, and 100 steps.

In Suppl. Figure 10 and 11 we show the mean absolute error (MAE):

$$MAE = \frac{1}{n} \sum_{i}^{n} |y_i - \hat{y}_i| \qquad \text{(Eq. 10)}$$

where $n$ is the number of samples, $y_i$ is the predicted value, and $\hat{y}_i$ is the true value for the Hurst exponent network (Suppl. Figure 2), and the Diffusion Coefficient Network (Suppl. Figure 5).

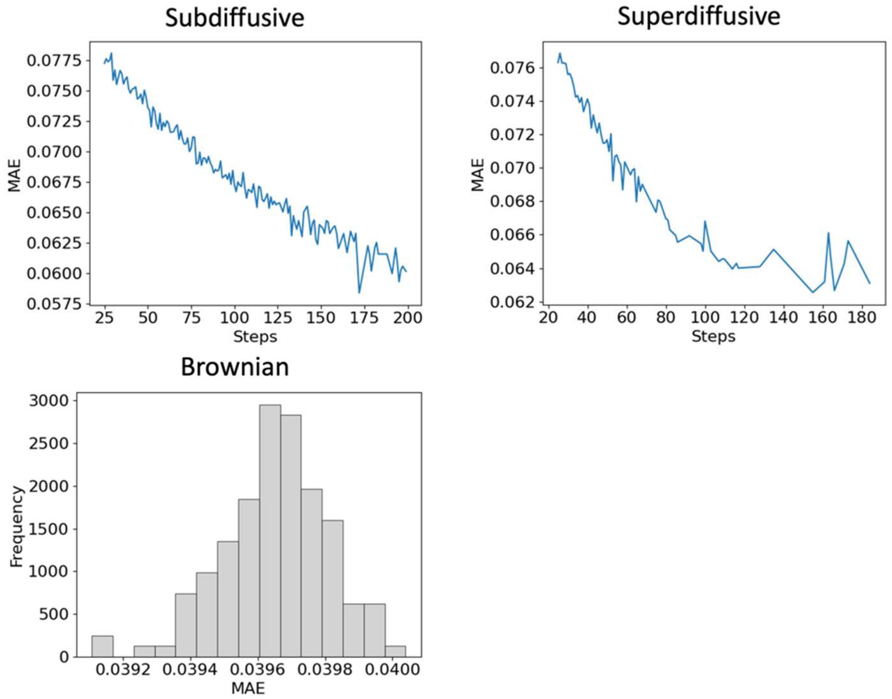


Suppl. Figure 10. Mean absolute error (MAE) for the 3 variants of the Hurst Exponent Network (Suppl. Figure 2) (subdiffusive, Brownian, and superdiffusive). In the case of the Brownian variant of the Hurst Exponent Network, the distribution of the MAE of our trained models is shown because there is no correlation between the steps and the MAE. This lack of correlation may be influenced by the localization error of the trajectories.

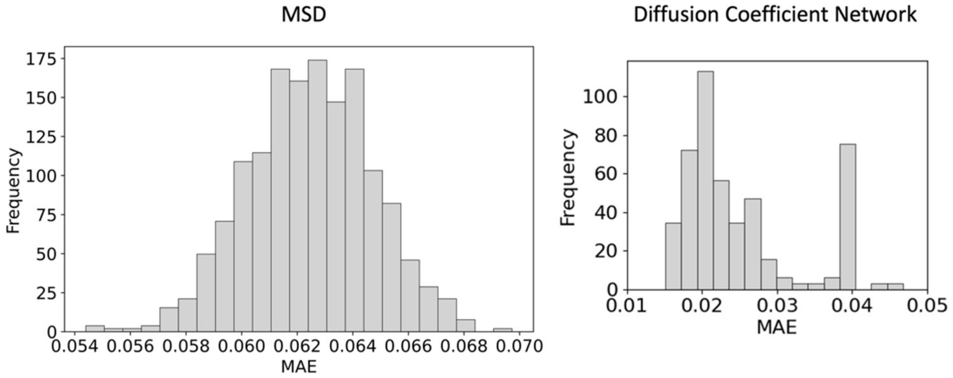


Suppl. Figure 11. Mean absolute error (MAE) of the estimation of the diffusion coefficient using a classical approach (MSD) (*left*) vs the estimation using the Diffusion Coefficient Network (Suppl. Figure

5) (*right*). The accuracy of the network approach clearly outperforms of the classical method (mean squared displacement, MSD).
